# Ezrin enrichment on curved membranes requires a specific conformation or interaction with a curvature-sensitive partner

**DOI:** 10.1101/297895

**Authors:** Feng-Ching Tsai, Aurélie Bertin, Hugo Bousquet, John Manzi, Yosuke Senju, Meng-Chen Tsai, Laura Picas, Stéphanie Miserey-Lenkei, Pekka Lappalainen, Emmanuel Lemichez, Evelyne Coudrier, Patricia Bassereau

## Abstract

One challenge in current cell biology is to decipher the biophysical mechanisms governing protein enrichment on curved membranes and the resulting membrane deformation. The ERM protein ezrin is abundant and associated with cellular membranes that are flat or with positive or negative curvatures. Using *in vitro* and cell biology approaches, we assess mechanisms of ezrin’s enrichment on curved membranes. We evidence that ezrin (ezrinWT) and its phosphomimetic mutant T567D (ezrinTD) do not deform membranes but self-assemble anti-parallelly, zipping adjacent membranes. EzrinTD’s specific conformation reduces intermolecular ezrin interactions, allows binding to actin filaments, and promotes ezrin binding to positively curved membranes. While neither ezrinTD nor ezrinWT senses negative membrane curvature alone, we demonstrate that interacting with curvature sensors I-BAR-domain proteins facilitates ezrin enrichment in negatively curved membrane protrusions. Overall, our work reveals new mechanisms, specific conformation or binding to a curvature sensor partner, for targeting curvature insensitive proteins to curved membranes.

## Introduction

Attachment of the plasma membrane or the membrane of organelles to the actin cytoskeleton network is an important feature that controls cell shape and function. Ezrin, a member of the ezrin-radixin-moesin (ERM) protein family, plays a crucial role in linking the actin cytoskeleton to membranes. Ezrin is involved in various physiological processes including cell migration, cell signaling and the establishment of cell polarity (McClatchey and Fehon, 2009) (Fehon et al., 2010) (Arpin et al., 2011). Ezrin consists of an N-terminal FERM (band 4.1, ezrin, radixin, moesin) domain that binds to phosphatidylinositol 4,5-bisphosphate (PIP_2_) lipids and membrane-associated binding partners, a structurally uncharacterized α-helical domain, and a C-terminal ERM association domain (C-ERMAD) that interacts with the FERM domain and the actin cytoskeleton. The head-to-tail intramolecular interaction between the FERM domain and the C-ERMAD keeps ezrin in a closed configuration where the actin binding site is masked (Bretscher et al., 1995). “Opening up” of ezrin requires the FERM domain to bind to PIP_2_, followed by the phosphorylation of the C-ERMAD (Fievet et al., 2004) (Pelaseyed et al., 2017). So far, the conformations of non-phosphorylated ezrin and of its phosphomimetic mutant have been studied either in solution, with or without PIP_2_ micelles (Pelaseyed et al., 2017) (Jayasundar et al., 2012), or on bare solid substrates (Liu et al., 2007). A recent paper reports a conformational change of the pseudophosphorylated mutant of ezrin upon binding to PIP_2_-containing supported bilayers (Shabardina et al., 2016), but a detailed analysis at high resolution at the nanometer scale of the different membrane-bound conformations of ezrin is still missing.

In cells, ezrin is one of the most abundant proteins with a concentration similar to that of actin (Viswanatha et al., 2013). Given the actin-membrane linking function of ezrin and the mechanical action of actin on membranes, it is essential for cells to precisely regulate the membrane localization of ezrin. Ezrin is enriched in actin-rich plasma membrane structures such as microvilli (Sauvanet et al., 2015), filopodia (Osawa et al., 2009) and at the edge of bacterial toxin-induced transendothelial cell tunnels (Stefani et al., 2017). In mutant mice lacking ezrin, enterocytes show thicker, shorter and misoriented microvilli, suggesting that ezrin contributes to microvilli morphology (Casaletto et al., 2011). In these plasma membrane protrusions ezrin is located at the cytosolic side wherein the membrane has a negative mean curvature. Therefore, this suggests ezrin has a strong affinity for negatively curved membranes; in other words, ezrin may be a negative membrane curvature-sensing protein. However, ezrin is also associated with some intracellular vesicles including endosomes (Chirivino et al., 2011), wherein the membranes have a positive mean membrane curvature. Moreover, ezrin is also found at flat regions of the plasma membrane, such as at the cortex-membrane interface and at the surface of membrane blebs (Charras et al., 2006). How the same protein can be a positive and a negative membrane curvature sensor is a conundrum that remains to be deciphered.

The complexity of studying ezrin-membrane interaction *in cellulo*, due to the presence of actin and cellular organelles, can be circumvented by using purified proteins and model membranes with controlled membrane curvature. In this study we combine cryo-electron microscopy (cryo-EM) and mechanical measurements using model membranes with cell biology approaches to compare wild type ezrin (ezrinWT) and its phosphomimetic mutant T567D (ezrinTD) for correlating ezrin conformation with its association to curved membranes. We show that: 1) both ezrinWT and ezrinTD assemble in an anti-parallel manner to tether adjacent membranes; 2) only the phosphomimetic mutant senses positive membrane curvature, likely due to its different conformation compared to ezrinWT; and 3) neither ezrinWT nor ezrinTD senses negative membrane curvature. Although *in vivo* most of ezrin is associated with membrane protrusions having negative membrane curvature, we show that the enrichment of ezrin and its phosphomimetic mutant on negatively curved membranes is facilitated by their direct interaction with curvature-sensing proteins, e.g. inverse-Bin-Amphiphysin-RVS (I-BAR) domain proteins. Altogether our data reveals new mechanisms for enriching ezrin on curved membranes, and reinforces the view of ezrin as a membrane-cytoskeleton linker and a scaffolding protein rather than a membrane shaper.

## Results

### Conformations of ezrin bound to PIP_2_-containing membranes revealed at the nanometer scale

To assess how the phosphorylation influences ezrin conformation and its binding to PIP_2_ membranes, we purified recombinant wild type ezrin with a histidine (His) tag (His-ezrinWT) and a phosphomimetic mutant where the threonine at position 567 was replaced by an aspartate (His-ezrinTD), mimicking the open configuration of ezrin (Fig. S1A) (Fievet et al., 2004). After proteolysis of the His tag, ezrinWT and ezrinTD were labeled with Alexa dyes for detection by confocal fluorescence microscopy (Fig. S1B). To measure ezrinWT or ezrinTD binding to membranes independently of membrane curvature, we used giant unilamellar vesicles (GUVs) having diameters of around 5 μm or more (thus flat at the scale of ezrin molecules) consisting of brain total lipid extract with or without 5 mole% PIP_2_. We measured the fluorescence signals of the labeled ezrin on GUVs by two independent techniques, confocal microscopy and flow cytometry. In agreement with previous reports, we found that ezrinTD and ezrinWT do not bind to GUVs lacking PIP_2_ (Fig. S1 C-F) (Carvalho et al., 2008). When bound to PIP_2_-containing GUVs, we observed homogeneous ezrin fluorescence signals on the membranes for both ezrinTD and ezrinWT (Fig. S1G top). Ezrin-decorated GUVs were globally spherical without optically detectable membrane deformation at bulk ezrin concentrations ranging from 20 nM to 4 μM. TO compare the binding affinities of ezrinTD and ezrinWT, we measured the absolute membrane surface fraction of fluorescent ezrin on GUV membranes, *Φ_v_*, at various bulk ezrin concentrations, *C_bulk_* (Fig. S1G bottom) (Sorre et al., 2012). By assuming a non-cooperative binding reaction, we fitted the binding curves with a hyperbola, *Φ_v_* = *Φ_max_* × *C_bulk_*/(*C_buik_* + *K_d_*), where *Φ_max_* is the maximum membrane surface fraction of ezrin and *K_d_* is the dissociation constant (Pollard, 2010). For *Φ_max_* = 12%, *K_d_* is equal to 1.2 μM and 4.2 μM for ezrinTD and ezrinWT, respectively. The estimated *K_d_* is comparable to the previously reported value obtained using similar free-standing membranes, large unilamellar vesicles (LUVs) (Blin et al., 2008), but are lower than reported values using solid-supported lipid bilayers (SLBs) (Bosk et al., 2011). Nonetheless, ezrinTD has a higher binding affinity for PIP_2_ than ezrinWT, as previously reported (Fritzsche et al., 2014) (Zhu et al., 2007), confirming the phosphomimetic substitution of T567 by an aspartic acid facilitates the binding of ezrin to PIP_2_.

We then investigated how ezrin organizes on PIP_2_-containing membranes at the nanometric level and in an aqueous ionic environment by using LUVs combined with cryo-EM. We prepared LUVs with diameters ranging between 100 and 500 nm by using the detergent removal method (Rigaud et al., 1998). In the absence of ezrin, LUVs were spherical and unilamellar (Fig. S2A). In the presence of ezrinTD or ezrinWT, to our surprise, we observed regular stacks of membranes tethered together by ezrinTD or ezrinWT (Fig. 1A). Three-dimensional reconstructions from cryo-tomography revealed that the lipid stacks are plate-like (Video S1 and S2). To enhance the signal-to-noise ratio, we performed two-dimensional (2D) single particle analysis by selecting pieces of stacks comprising two bilayers and the protein material between them. The class-averages revealed densely packed globular domains on the lipid layers that are 4.5 nm apart from the membrane (as measured from the center of the globular domains to the closest lipid leaflet) (Fig. 1B). Further 2D image analysis centered on the globular domains revealed their dimensions to be 4 nm long by 3 nm wide for both ezrinTD and ezrinWT (Fig. 1C). This is in good agreement with the reported FERM domain structure of ezrin (Smith et al., 2003), suggesting that these globular domains correspond to the FERM domain. The α-helical domain and C-ERMAD were less resolved under these experimental conditions. Nevertheless, we found that the distances measured between the centers of the globular domains of the two opposing molecules sandwiched between the lipid layers were different: 24.1 ± 1.3 nm for ezrinTD and 28.7 ±1.2 nm for ezrinWT (error bar represents standard deviation) (Fig. 1D). These distances are comparable with the proposed model of the fully open ERM proteins (24-32 nm) (Fehon et al., 2010) (Jayasundar et al., 2012) (Phang et al., 2016), and with lengths measured for ezrin (15-45 nm) (Liu et al., 2007) in its phosphorylated/open states and in the absence of PIP_2_. Thus, our observations suggest that on the tethered membranes, both ezrinTD and ezrinWT have a different but open conformation. By measuring the frequency of membrane tethering at the same protein bulk concentration, we observed that ezrinTD induced more membrane tethering as compared to ezrinWT (Fig. 1E). Taken together, our results demonstrate that on PIP_2_-containing membranes, both ezrinWT and ezrinTD self-assemble into brush-like pairs that dramatically disrupt and reorganize LUVs into bilayer stacks. Moreover, ezrinTD and ezrinWT molecules exhibit different lengths that reflect their distinct conformations.

**Figure 1.**
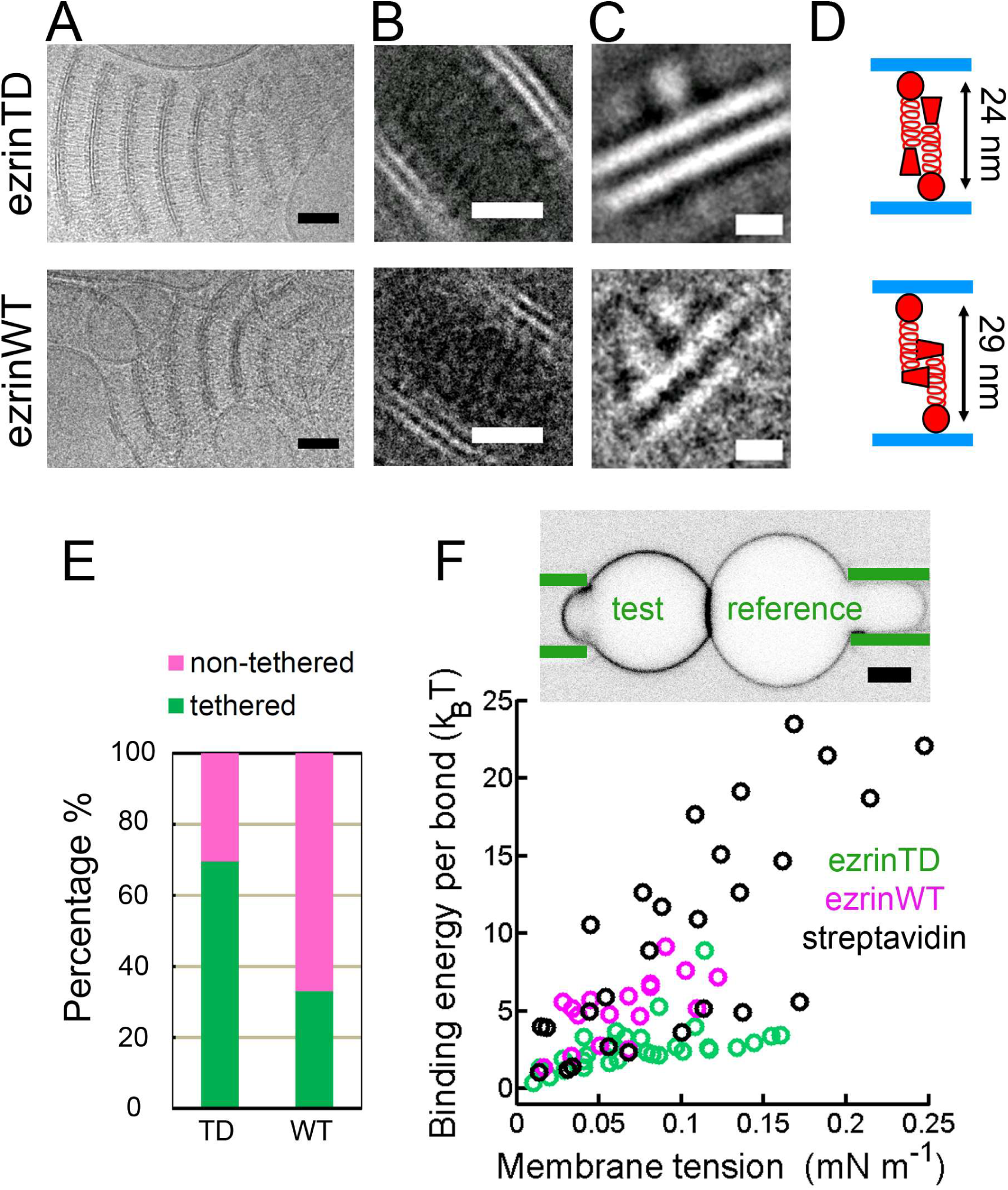
EzrinTD and ezrinWT have distinct conformations on PIP_2_-containing membranes. (A) Representative cryo-electron micrographs of PIP_2_-membranes tethered by ezrinTD (0.2 μM) or ezrinWT (1.2 μM). Scale bars: 50 nm. (B-C) Representative two-dimensional class averages of ezrinTD- and ezrinWT-tethered membranes. The distances between the globular domains of ezrinTD and ezrinWT between two tethered membranes are 24.1 ± 1.3 nm and 28.7 ± 1.2 nm, respectively (see statistic details in the Materials and Methods). Scale bars: (B) 20 nm and (C) 5 nm. (D) Cartoons illustrating ezrin-membrane tethering by ezrinTD and ezrinWT. (E) Percentages of tethered and non-tethered membranes deduced from cryo-EM. EzrinTD andezrinWT bulk concentration: 1.2 μM. Total membrane length measured: 94728 nm (ezrinTD)and 77115 (ezrinWT) for n = 3 experiments. Measurements were taken from distinct samples. *p* < 0.0001 (z-test), comparing the percentage of tethering of ezrinTD and ezrinWT. (F) Dual micropipette assay (Top) Representative confocal image of the test GUV and the reference GUV tethered by ezrinTD (inverted grayscale). The green lines indicate the micropipettes. Scale bar, 5 μm (Bottom) Membrane tethering strengths of ezrinTD (N = 8 GUVs, n = 4 experiments), ezrinWT (N = 4 GUVs, n = 3 experiments) and streptavidin bonds (N = 5 GUVs, n = 2 experiments). The following figure supplements are available for figure 1: **Figure supplement 1**. Analysis of purified recombinant ezrinTD and ezrinWT, and their binding to PIP_2_-containing membranes. **Figure supplement 2B**. Representative cryo-electron micrograph of LUVs. **Figure supplement 3**. Dual micropipette experiments with GUVs tethered by ezrinTD or ezrin. **Video supplement 1**. Cryo-tomography of ezrinTD tethered membrane stacks. **Video supplement 2**. Cryo-tomography of ezrinTD tethered membrane stacks.

If ezrin and its phosphomimetic mutant have different conformations on membranes, the binding energy between two tethered membranes should be different. To test this hypothesis, we performed experiments using a dual micropipette aspiration assay. Two GUVs decorated with ezrin were brought in contact while controlling the membrane tension of both GUVs by the micropipette aspiration pressure (Evans and Needham, 1988) (Noppl-Simson and Needham, 1996). We followed the previously established experimental procedure in which one GUV, named the reference GUV, was under high tension to ensure a spherical shape while adjusting the membrane tension of the other GUV, named the test GUV (Franke et al., 2006). At low membrane tension of the test GUV, we observed an enrichment of ezrin fluorescence signal in the contact zone while the test GUV had a lemon-like shape, indicating ezrin tethering (Fig. 1F and S3A). No membrane tethering was observed when ezrin was absent. Monitoring ezrin mobility in the tethering zone with fluorescent recovery after photobleaching (FRAP) revealed that only 10% of ezrinTD and ezrinWT diffuse freely at the interface (Fig. S3B), confirming the formation of a stable contact zone resulting from ezrin tethering. We deduced the tethering energy per ezrin pair bond between the two GUVs by using their contacting geometry at a given membrane tension (see Fig. S3C for the schematic and Methods for details) (Franke et al., 2006). We compared ezrin tethering energy to that of the streptavidin-biotin bond as a reference. For both the ezrin-ezrin bond and the biotin-streptavidin bond, the binding energy was dependent on the membrane tension (Fig. 1F). This tension dependence may arise from the electrostatic repulsion of the opposing membranes, as suggested previously (Franke et al., 2006). Nonetheless, at all tensions probed the tethering energy calculated for ezrinWT was on average higher than that of ezrinTD but lower than that of the biotin-streptavidin bond (Fig. 1F) (Noppl-Simson and Needham, 1996). Thus, when associated to membranes the different conformations of ezrinWT and ezrinTD lead to different strengths of the intermolecular interaction of ezrinWT and ezrinTD, as revealed by their different tethering energy.

### Brush-like ezrinTD assembly connects F-actin and membranes

Extensive experimental evidence points out that the C-ERMAD of ERM proteins has to be phosphorylated for their association with F-actin (Fievet et al., 2004) (Fritzsche et al., 2014) (Nakamura et al., 1999). However, studies using SLBs have proposed that the binding of ezrinWT to PIP_2_ is sufficient to change its conformation and to expose its actin binding site (Bosk et al., 2011). The fact that ezrinWT and ezrinTD display different conformations on membranes motivated us to examine whether ezrinTD and ezrinWT bridge F-actin to PIP_2_-containing membranes. We first assessed actin recruitment on GUVs. When GUV membranes were covered with AX546-labeled ezrinTD, the recruitment of AX488 phalloidin-labeled muscle F-actin to the membranes was clearly visible (Fig. 2A), as F-actin formed a dense, network-like structure on the GUVs (Fig. S4A). In contrast, no detectable muscle F-actin was present on AX546-labeled ezrinWT-coated GUVs (Fig. 2B), while F-actin was clearly visible in the background as in the control experiment where there was no ezrin on GUVs (Fig. 2 C and F). We obtained the same results when replacing muscle F-actin with non-muscle F-actin (Fig. 2 D and E, and Fig. S4B). Our observation seems inconsistent with a previous study using SLBs on silicon substrates where F-actin recruitment on ezrinWT decorated PIP_2_-membranes was observed (Bosk et al., 2011). However, in Bosk *et al*., it was reported that ezrinWT formed large immobilized clusters on SLBs even at low PIP_2_ density that might be attributed to ezrinWT activation. In contrast, we never observed clusters of ezrinTD or ezrinWT on free-standing GUV membranes. Moreover, a strong enhancement for F-actin recruitment was also observed by Bosk *et al*. in the case of ezrinTD as compared to ezrinWT. Here, our observations clearly demonstrate that PIP_2_ is not sufficient for the recruitment of F-actin by ezrinWT at the surface of GUVs, which might be closer to the biological situation than SLBs.

**Figure 2.**
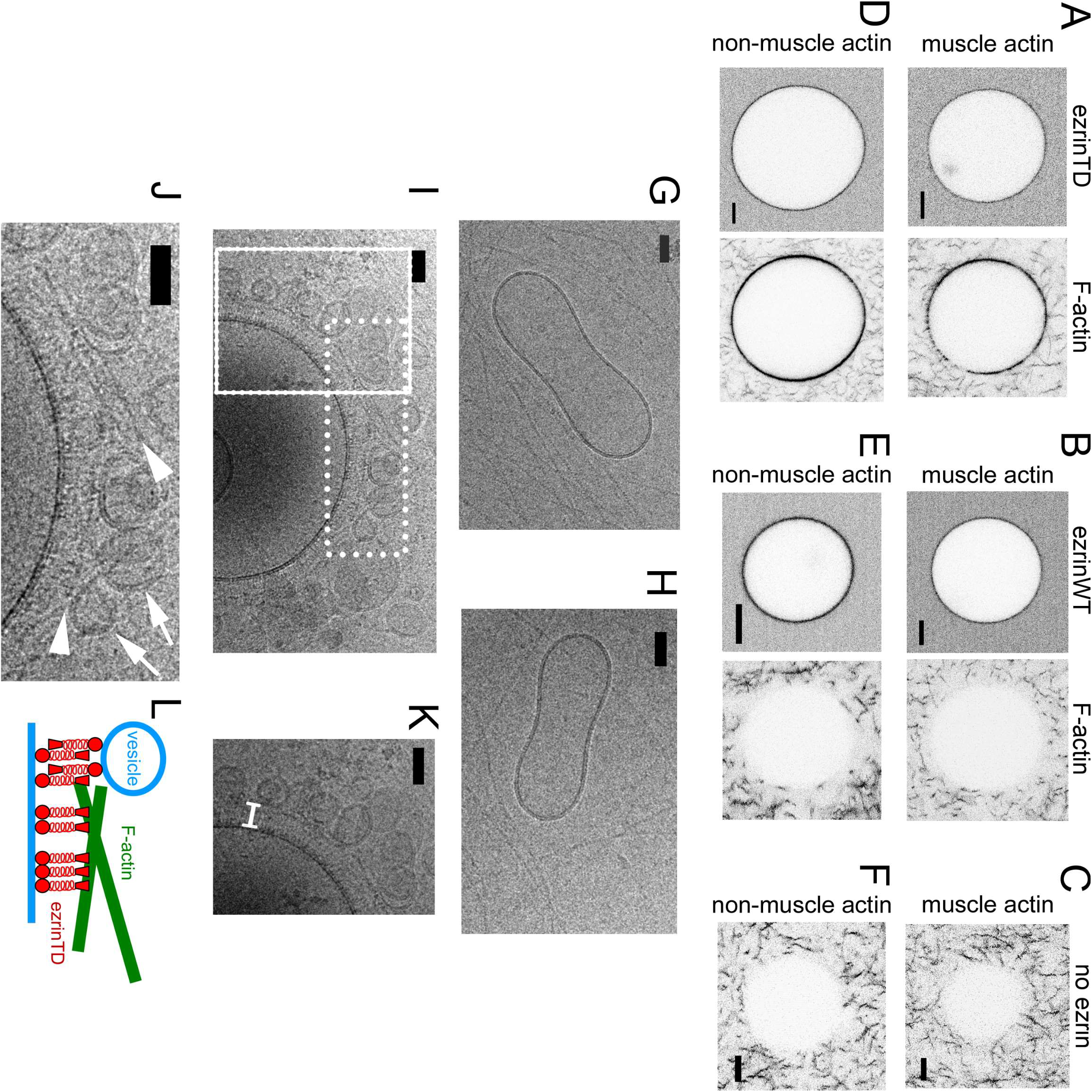
EzrinTD forming a brush-like structure bridges F-actin and the membrane. (A-F) Representative confocal images of GUVs coated with ezrinTD (A and D), ezrinWT (B and E) or in the absence of ezrin as a control (C and F), in the presence of muscle (A, B and C) or non-muscle (D, E and F) F-actin. The bulk concentrations of muscle and non-muscle F-actin are 0.4 and 0.6 μM, respectively, and the bulk concentrations of ezrinTD and ezrinWT are both 0.5 μM. Scale bars, 5 μm. Inverted grayscale images were shown. (G-J) Representative cryo-electron micrographs of PIP_2_-containing LUVs incubated with muscle F-actin in the absence (G and H) and presence (I-K) of ezrinTD. (J and K) Enlargements of the regions marked by the dotted and solid line rectangle in (I), respectively. In (J) Arrowheads indicate F-actin and arrows indicate tethered vesicles. In (K) white bracket indicates brush-like ezrinTD assembly. EzrinTD and F-actin concentrations are 0.3 μM and 1 μM, respectively. Scale bars, 50 nm. (L)Cartoon illustrating brush-like ezrinTD assembly recruiting F-actin and tethering vesicles to membranes. The following figure supplements are available for figure 2: **Figure supplement 4**. Network-like F-actin organization on ezrinTD-decorated GUV membranes. **Figure supplement 5**. F-actin on vesicle surfaces decorated with ezrinTD.

To resolve the membrane-ezrinTD-actin organization at nanometric resolution, we carried out cryo-EM experiments using LUVs. In the absence of ezrinTD, no interaction between LUVs and muscle F-actin was observed (Fig. 2 G and H). In the presence of ezrinTD, we observed that actin filaments were parallel to the membrane surface, following the contour of the membrane, (Fig. 2 I and J arrowheads, and Fig. S5) with a brush-like assembly of ezrinTD bridging F-actin and the membrane (Fig. 2 I and K). The distance between actin filaments and the membrane is about 25 nm, similar to the length of the open ezrinTD molecule (Fig. 1D). Moreover, small vesicles were also found in contact with the ezrinTD brush (Fig. 2 I and J arrows). Our results thus confirm that a negative charge at position T567 of C-ERMAD, i.e. mimicking ezrin phosphorylation, mediates a conformational change of ezrin that then makes ezrin amenable to link F-actin to membranes. Moreover, we reveal that on PIP_2_-containing membranes ezrinTD sits perpendicularly to the membrane with its FERM domain interacting with PIP_2_ and its phosphomimetic C-ERMAD domain protruding from the membrane to engage interactions with actin filaments (Fig. 2L). This is in contrast to the previous hypothesis suggested the configuration of ezrin dimers on membranes by structural analysis in solution (Jayasundar et al., 2012) (Phang et al., 2016).

### Phosphomimetic mutation enables the positive membrane curvature sensing of ezrin

Since ezrinWT and ezrinTD have different conformations when associated with quasi-flat PIP_2_-containing LUVs and GUVs, we next investigated how they interact with highly curved galactocerebroside nanotubes (Dang et al., 2005). These rigid nanotubes with a uniform external diameter of 25 nm were composed of galactocerebrosides supplemented with 15 mole% EPC and 5 mole% PIP_2_. Thus these nanotubes offer a suitable approach to probe positive curvature sensing of proteins by cryo-EM. They were dispersed in solution in the absence of protein (Fig. S2B). Addition of ezrinWT or ezrinTD promoted different types of nanotube assemblies. In the presence of ezrinTD, 68.5% of the tubes formed disordered assemblies (henceforth named disordered) and 25% were isolated nanotubes (henceforth named isolated) (Fig. 3 A-C). In the presence of ezrinWT, we observed up to 42% of isolated nanotubes and instead of a disordered organization, 58% of the tubes formed parallel stacks (henceforth named stacked) (Fig. 3 A-C). These stacks have a regular spacing of 25.8 ± 2.4 nm between the centers of the opposing FERM domains, similar to the distance between the tethered bilayers observed in the LUV experiments (Fig. 1D). The difference in the nanotube assemblies between ezrinWT and ezrinTD suggests a difference in the interaction and organization of individual molecules with the nanotubes. In the presence of ezrinTD all isolated tubes (total 15 tubes) were covered with proteins. Moreover, we observed that on tubes, some ezrinTD proteins bound along the short axis of the tube, with their whole length in contact with the tube surface (Fig. 3B top, arrows and Fig. S6A), while the rest was attached perpendicularly (Fig. 3B top, arrowheads and Fig. S6B). In contrast, in the presence of ezrinWT, only 55% (total 26 tubes) of the isolated tubes were decorated with proteins, showing a weaker affinity for positively curved membranes compared to that of ezrinTD. Moreover, we observed that ezrinWT proteins bound perpendicularly to the nanotube surface (Fig. 3B bottom, arrowheads). Altogether, these observations indicate that ezrinTD and ezrinWT can tether membranes with strong positive curvature but they induce different types of organization due to their different conformations.

**Figure 3.**
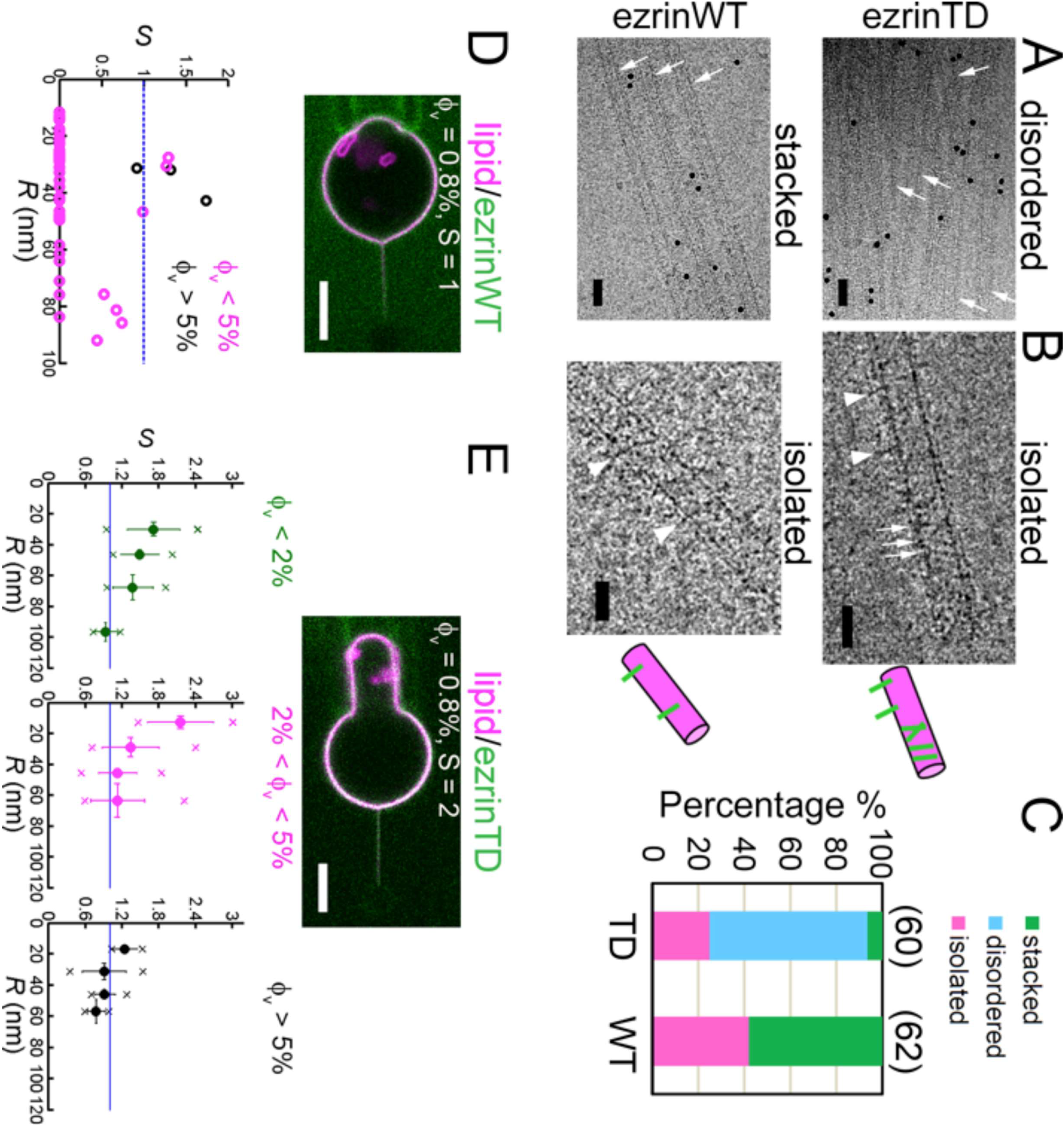
EzrinTD binds to positively curved membranes. (A) Rigid lipid nanotubes assembled by ezrin: representative cryo-electron micrographs of ezrinTD inducing randomly-oriented nanotube assembly (disordered) and ezrinWT inducing parallel stacks (stacked). Arrows indicate some of the nanotubes in both cases. Scale bars: 50 nm. The black dots are gold particles. (B) Isolated nanotubes in the presence of ezrinTD or ezrinWT. Arrows and arrowheads indicate proteins on the nanotubes. Scale bars: 25 nm. Protein concentrations: 1.2 μM for both ezrinTD and ezrinWT. Nanotube concentration: 0.1 g.L^−1^. Cartoons illustrate isolated nanotubes decorated by ezrinTD and by ezrinWT in the corresponding EM images. (C) Percentages of membrane nanotubes being disordered, forming stacks or being isolated in the presence of ezrinTD or ezrinWT, deduced from cryo-EM. Sample numbers are indicated in brackets. n = 2 experiments per condition. Measurements were taken from distinct samples. Numbers of tubes measured are N = 60 and N = 62 for ezrinTD and ezrinWT, respectively. Statistic test (chi-square test): *p* ≈ 1.5 × 10^−9^ for stacked tubes and *p* = 0.0477 for isolated tubes. (D and E) Nanotube pulling assay to probe ezrin positive membrane curvature sensing. (Top) Representative confocal images of ezrinWT (D) and ezrinTD (E) binding on a membrane nanotube pulled from a GUV aspirated in a micropipette. The nanotube is held by a bead (not fluorescent) that is trapped by optical tweezers. Scale bars, 5 μm. (Bottom) Sorting ratio, *S*, as a function of tube radius, *R*, for ezrinWT (D) and ezrinTD (E). ϕ_v_: ezrin membrane surface fraction. Blue lines indicate *S* = 1. In (D), measurements were collected from N = 13 GUVs, n > 3 independent experiments. In (E), measurements were collected from N = 63 GUVs, n > 3 independent experiments. Error bars indicate standard deviations, circles are mean and X symbols are maximum and minimum for each condition, excluding *S* = 0 data. The following figure supplements are available for figure 3: **Figure supplement S2B**. Representative cryo-electron micrograph of galactocerebroside nanotubes in the absence of ezrin. **Figure supplement S6**. EzrinTD on isolated nanotubes. **Figure supplement S7**. Binding of ezrin on GUVs, positive curvature-induced sorting of PIP_2_, ezrinTD and the FERM domain of ezrin on membrane nanotubes.

The different conformations observed between ezrinWT and ezrinTD on rigid galactocerebroside nanotubes suggest a different ability to sense positive membrane curvature. We thus quantified the positive curvature sensing ability of ezrin by measuring the enrichment of ezrinWT and ezrinTD on positively curved membranes relative to flat ones. To this end, we performed tube pulling experiments where a fluid membrane nanotube was pulled outwards from a micropipette-held GUV while injecting ezrin next to the GUV with another micropipette, as previously done with some curvature sensing proteins (Sorre et al., 2012). At a given membrane tension, after injecting AX488 labeled ezrinWT or ezrinTD for 2-3 mins, the membrane coverage of ezrin on GUVs generally plateaued (Fig. S7A). By controlling the aspiration pressure, the membrane tension of a GUV can be adjusted. In the absence of protein binding, the radius of the nanotube *R* is set by the membrane tension σ and the membrane bending rigidity κ of the GUV, 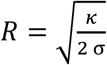 (Derenyi et al., 2002). We therefore have a direct control of the nanotube radius, typically ranging from 10 nm to 100 nm. By calculating the ratio of lipid fluorescence intensities in the tube and in the GUV, we can derive the tube radius (see Methods for details). Our experimental system allows us to assess curvature sensing by measuring the ratio of ezrin fluorescence intensities on the tube (positively curved membranes) and on the GUV (flat membranes), normalized by the ratio of lipid fluorescence intensities on both structures to calculate the sorting ratio *S*, 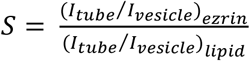 (Sorre et al., 2012). For experiments where the fluorescence signals of ezrin on tubes were too low to be measured (close to the noise level), we set *S* equal to 0. We first checked that in the absence of ezrin, no sorting of PIP_2_ in membrane tubes was observed (Fig. S7B), validating that protein sorting will result from the property of the protein only. No binding on tubes was detected (*S* = 0) for ezrinWT, in a relatively large fraction of experiments; when detectable, ezrinWT was not enriched on tubes as demonstrated by *S* ≅ 1 (Fig. 3D). In contrast, binding and enrichment on tubes was detected for ezrinTD. We grouped the sorting data into three classes corresponding to different membrane surface fractions of ezrinTD on the GUVs, *Φ_v_*, since curvature-induced sorting is expected to depend on *Φ_v_* (Sorre et al., 2012) (Zhu et al., 2012) (Prevost et al., 2015). We observed that *S* monotonically increased when the tube radius decreased and *S* was larger than 1, showing that ezrinTD is a positive curvature sensor (Fig. 3E and Fig. S7C). As expected (Sorre et al., 2012) (Zhu et al., 2012) (Prevost et al., 2015), *S* was lower for the highest *Φ_v_* values (Fig. 3E and Fig. S7C). However, the positive curvature sensing ability of ezrinTD is weaker than previously reported curvature-sensing proteins (Sorre et al., 2012) (Zhu et al., 2012) (Prevost et al., 2015) as *S* is lower than 3 at low protein membrane coverage. We then examined if the FERM domain of ezrin senses positive membrane curvature by performing the same tube pulling experiments as for ezrin. The enrichment of the FERM domain on positively curved membranes was clearly observed and even stronger than that of ezrinTD, as sorting values up to 4 were observed (Fig. S7D). Thus, our results demonstrate that the conformation of ezrinTD allows its FERM domain to sense positive membrane curvature.

### Neither ezrinTD nor ezrinWT senses negative membrane curvature

We next wondered how ezrin is enriched into cellular protrusions wherein membranes have a negative mean curvature. To this end, we pulled membrane nanotubes from GUVs encapsulating ezrin and measured the sorting ratio *S*, as described earlier (Prevost et al., 2015). We observed that ezrin was present on the external leaflet during GUV preparation and could be effectively detached at high salt concentration 300 mM) (Fig. S8). We set *S* = 0 for experiments where the fluorescence signals of ezrin were too weak for detection. For both ezrinTD and ezrinWT, we observed no clear dependence of the sorting ratio *S* on tube radius within the experimental accessible tube radii. At low protein coverage on the GUVs (<5%) or higher (>5%) the sorting ratio *S* fluctuates around 1, indicating the absence of ezrinTD or ezrinWT enrichment in tubes (Fig. 4). Our results therefore demonstrate that ezrin is not a negative membrane curvature sensor.

**Figure 4.**
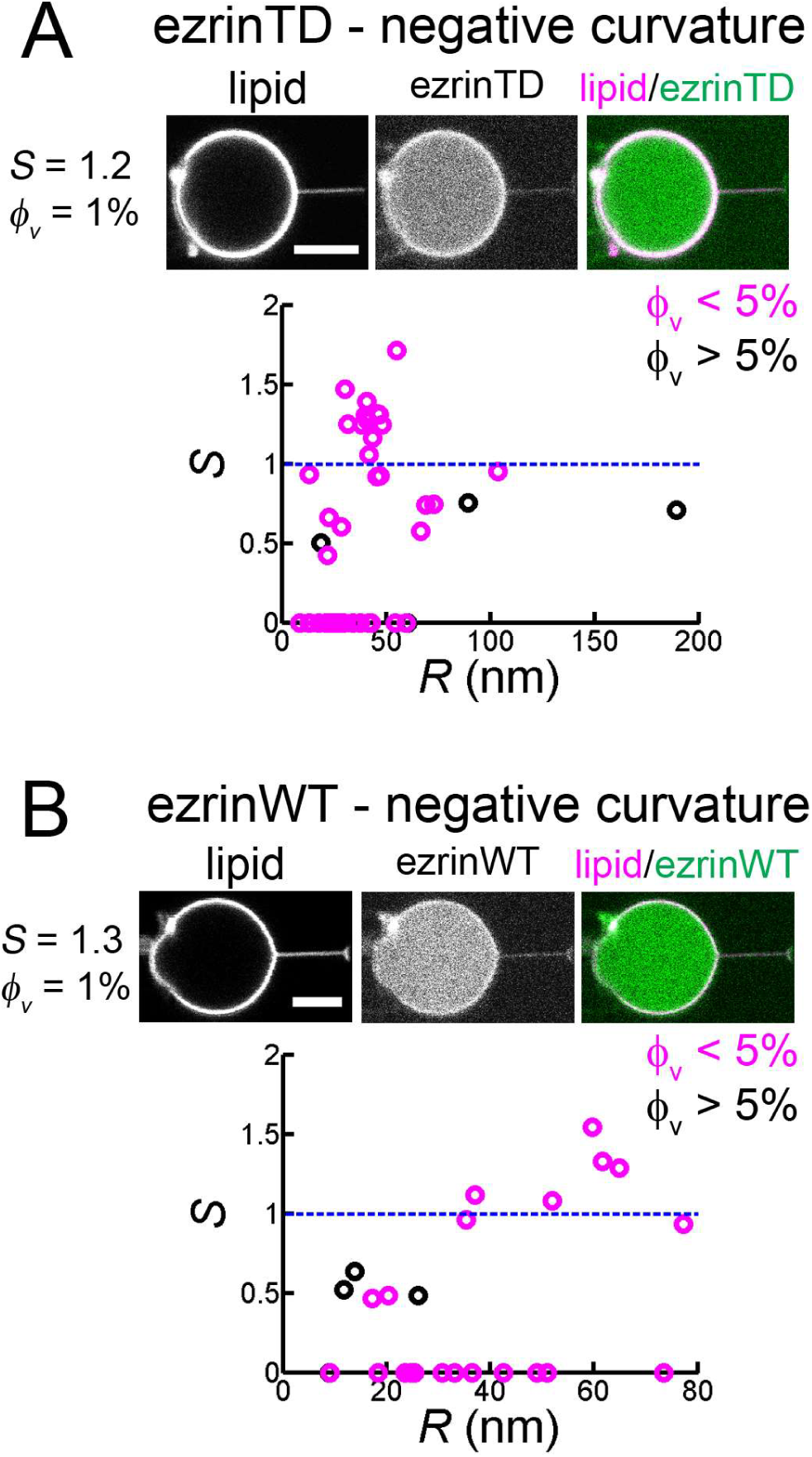
Both ezrinTD and ezrinWT do not sense negative membrane curvature. (A and B) Tube pulling experiments from GUVs encapsulating ezrinTD (A) or ezrin WT (B). (Top) Representative confocal images of ezrin sorting experiments, corresponding to (A) a sorting ratio *S* = 1.2 at *ϕ_v_*= 1% and (B) *S* = 1.3 at *ϕ_v_*= 1%, respectively. Scale bars, 5 μm. (Bottom) Sorting ratio, *S*, as a function of membrane tube radius *R* and for different ezrin surface fractions on the GUVs, ϕ_v_ Measurements were collected from N = 26 GUVs (n > 3 experiments), and N = 7 GUVs (n = 3 experiments), for ezrinTD and ezrinWT, respectively. Blue lines indicate *S* = 1. The following figure supplement is available for figure 4: **Figure supplement S8**. Salt-induced desorption of ezrinTD and ezrinWT bound to the outer leaflet of GUVs.

### EzrinWT and ezrinTD are enriched in negatively curved membranes by interacting with I-BAR domain proteins

Since ezrin has no intrinsic affinity for negatively curved membranes although it shows a cellular enrichment on negatively curved membranes, we investigated whether ezrin is recruited through partner proteins that are enriched in cellular protrusions. IRSp53, an I-BAR domain containing protein, was found to colocalize with ezrin in microvilli (Garbett et al., 2013). Moreover, IRSp53 is involved in the initiation of filopodia (Disanza et al., 2013), and is enriched *in vitro* inside model membrane nanotubes (Prevost et al., 2015). Therefore, IRSp53 is a potential candidate to recruit ezrin to membrane protrusions. Indeed, using super-resolution structured illumination microscopy, we observed that patchy fluorescent signals of GFP-IRSp53 or its GFP-I-BAR domain overlapped partially with the endogenous ezrin in cellular protrusions of LLC-PK1 epithelial cells (Fig. 5 A and B). We confirmed the association of ezrin with IRSp53 or with its I-BAR domain by co-immunoprecipitating the endogenous ezrin with the GFP-IRSp53 or GFP-I-BAR domain in HeLa cells (Fig. 5 C and D). GFP-IRSp53 and GFP-I-BAR immunoprecipitated ezrin to the same extent (Fig. 5D) regarding their expression level shown Fig. 5C. To assess whether ezrin and the I-BAR domain of IRSp53 interact directly, we performed a GUV-binding assay. We generated GUVs containing DOPC and DOGS-Ni-NTA lipids, without PIP_2_ (Ni-GUVs). In the absence of the I-BAR domain, ezrin does not bind to Ni-GUVs (Fig. 5E). We then incubated Ni-GUVs first with His-tagged I-BAR domain and then with ezrinWT or ezrin TD. In the presence of the His-tagged I-BAR domain, bound to Ni-GUVs via the His/Ni-NTA interaction, we found both ezrinTD and ezrinWT binding on Ni-GUVs (Fig. 5F). As a control, we checked that ezrin does not bind to Ni-GUVs covered with his-GFP (Fig. 5G). These observations demonstrate a direct interaction between the I-BAR domain and ezrinWT and ezrinTD.

**Figure 5.**
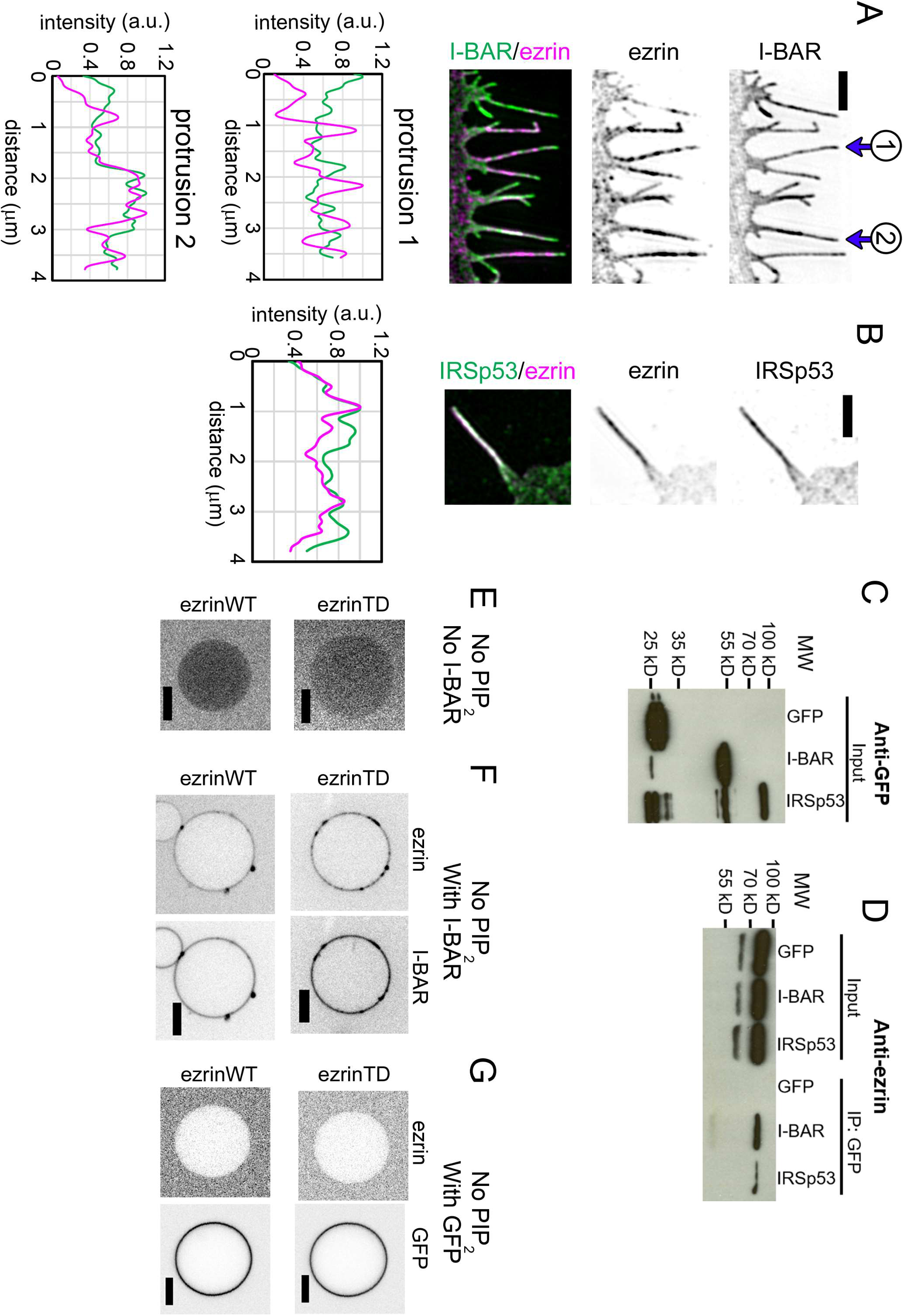
Ezrin partially colocalizes with IRSp53 in cellular protrusions and directly interacts with the I-BAR domain of IRSp53. (A and B) (Top) Representative structured illumination microscopy images of cellular protrusions of LLC-PK1 cells transfected with plasmids encoding GFP-I-BAR domain (A) or GFP-IRSp53 (B), and immunolabeled for endogenous ezrin. Scale bars, 2 μm. Inverted grayscale images were shown, unless color codes were indicated in the figure. (Bottom) Normalized fluorescence intensity profiles of protrusions indicated in (A) and the protrusion in (B). The distance zero is at the tip of the protrusions and the distance goes along the protrusions until the base of the protrusions at the cell edge. (C and D) GFP, GFP-IRSp53 or GFP-I-BAR domain transfected HeLa cells were lysed and immunoprecipitated (IP) using GFP-trap. Cell lysates (input) (C) and IP (D) were analyzed by Western blot for the expression of GFP, GFP-I-BAR domain and GFP-IRSp53 using an anti-GFP antibody (C) and pull down of endogenous ezrin using an anti-ezrin antibody (D). (E-G) Representative confocal images of ezrinTD or ezrinWT binding to Ni-GUVs in the absence of both PIP_2_ and the I-BAR domain (E), or in the absence of PIP_2_ but in the presence of the I-BAR domain (F), or in the absence of PIP_2_ but in the presence of GFP (G) observed in at least 3 independent experiments for (E) and (F), and in 2 independent experiments for (G). Arrowheads and arrows indicate clusters of ezrin and the I-BAR domain in (F). Protein bulk concentrations: for ezrinTD and ezrinWT, 1 μM, and for the I-BAR domain, 2 μM. Scale bars, 5 μm. Inverted grayscale images were shown.

We next examined whether ezrin is enriched on negatively curved membranes in the presence of the I-BAR domain of IRSp53. Encapsulation of two proteins in GUVs is very challenging, thus we took advantage of the spontaneous GUV membrane tubulation induced by the I-BAR domain (Saarikangas et al., 2009) (Mattila et al., 2007) (Barooji et al., 2016). Incubation of PIP_2_-containing GUVs with the I-BAR domain of IRSp53 led to membrane tubular invaginations towards the interior of the GUVs (Fig. S9A), as shown previously (Saarikangas et al., 2009) (Mattila et al., 2007) (Barooji et al., 2016). To validate our assay, we quantified the relative distribution of the I-BAR domain on GUVs and on the corresponding tubules by measuring the sorting ratio *S* of the I-BAR domain. In this tubulation assay, we cannot use any calibration method based on lipid fluorescence intensities to determine the tube radii. We thus used the membrane fluorescence ratio on the tube and on the corresponding GUV, (*I_tube_/I_vesicle_*)*_membrane_*, as a relative measurement of the tubule radius. We observed a non-monotonic enrichment of the I-BAR domain depending on the membrane fluorescence ratio (*I_tube_/I_vesicle_*)*_membrane_* (Fig. S9B), in close agreement with a previous report using the tube pulling assay (Prevost et al., 2015). Consistent with the previous report by Chen *et al*. (Chen et al., 2015), we observed I-BAR-induced tube formation at low I-BAR domain membrane coverage on GUVs (<2%). Moreover, in agreement with the previous study by Prévost *et al*. (Prevost et al., 2015), we observed a stronger sorting of the I-BAR domain at lower I-BAR domain membrane coverage on GUVs (<2%) than at higher membrane coverage (>5%) (Fig. S9B). By comparing our results with those obtained by Prévost *et al*. (Prevost et al., 2015), we could estimate tube radii considering that the maximum sorting of the I-BAR domain at (*I_tube_/I_vesicle_*)*_membrane_* ≅ 0.3 should correspond to the intrinsic spontaneous radius of the I-BAR domain, *R* ≈ 20 *nm* (Fig. S9B).

To assess if ezrin is enriched in the tubules induced by the I-BAR domain of IRSp53, GUVs were first incubated with ezrin followed by the addition of the I-BAR domain. We observed that the I-BAR domain deformed ezrin-coated membranes and that ezrinTD or ezrinWT was present on I-BAR domain-induced tubules (Fig. S10). The stronger fluorescence signal of ezrinTD or ezrinWT on I-BAR domain-induced tubules, as compared to those on GUVs, clearly indicates the enrichment of ezrin on the negatively curved tubular membranes (Fig. 6 A and B). Moreover, we observed a strong sorting of ezrinTD with a sorting ratio *S* up to 8 at the lowest ezrinTD membrane coverage (<2%), which decreased when *ϕ_v_* increased (Fig. 6C). The maximum sorting of ezrinTD at a membrane fluorescence ratio ≈ 0.4 corresponds to a tube radius of about 30 nm (Fig. 6C). Similarly, ezrinWT also enriched in I-BAR domain-induced tubules (Fig. 6D).

**Figure 6.**
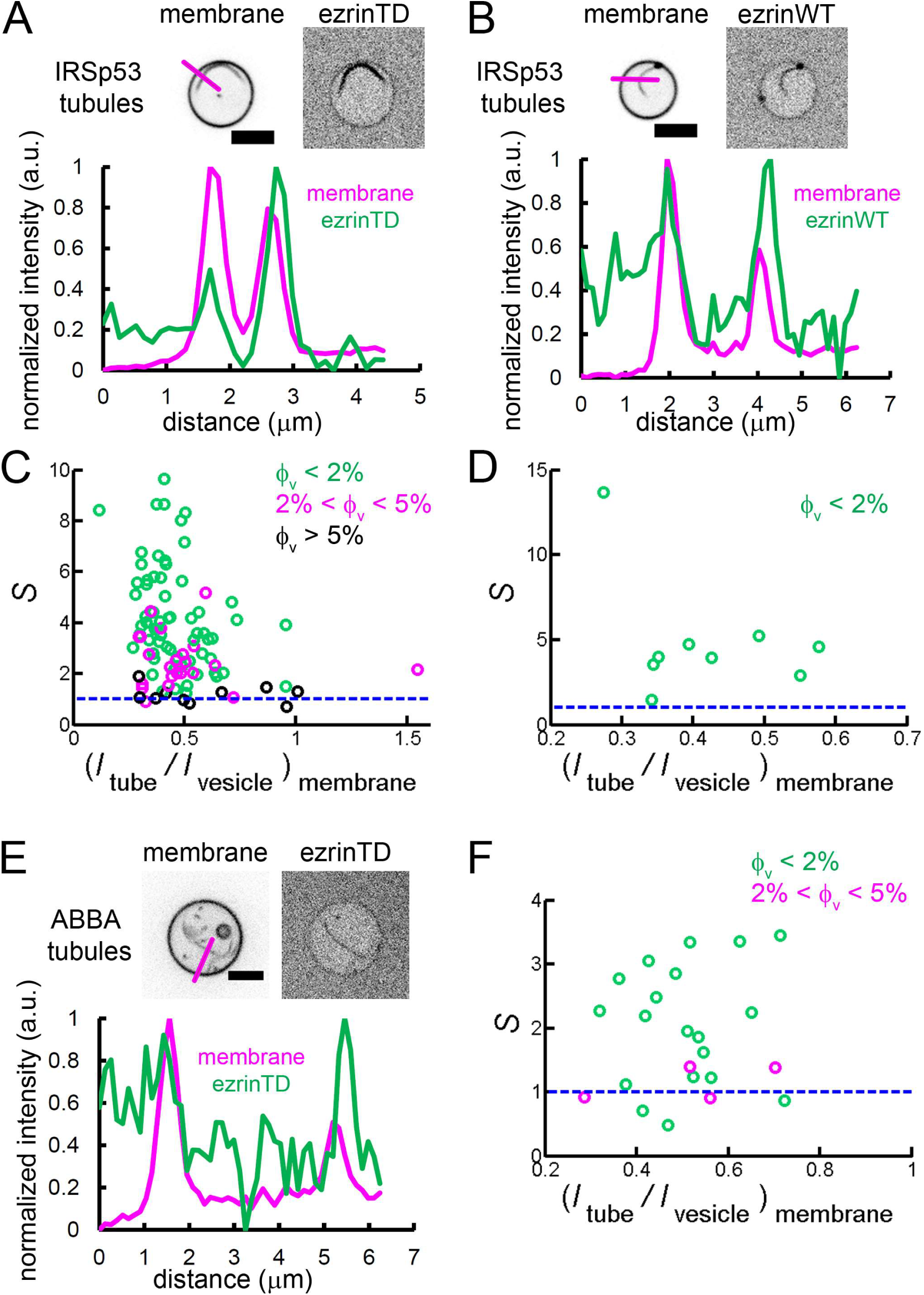
Enrichment of ezrinTD and ezrinWT in I-BAR domain induced tubules. (A and B) (Top) Representative confocal images of ezrinTD (A), and ezrinWT (B) in IRSp53I-BAR domain induced tubules. (Bottom) Normalized fluorescence intensity profiles along the line drawn from outside the GUV towards the interior of the GUV, as indicated in the top image. Inverted grayscale images were shown. (C and D) Sorting of ezrinTD (C) and ezrinWT (D) in IRSp53 I-BAR domain-induced membrane tubules at different membrane coverages of ezrin. (*I_tube_/I_vesicle_*)_*membrane*_ ≅ 0.4 corresponds to a tubule radius of about 30 nm. Scale bars, 5 μm. In (C), measurements were collected from N = 94 GUVs, n = 6 experiments. In (D), measurements were collected from N = 9 GUVs, n = 4 experiments. (E) (Top)Representative confocal images of ezrinTD in ABBA I-BAR domain-induced tubules. Inverted grayscale images were shown. (Bottom) Normalized fluorescence intensity profiles along the line drawn from outside the GUV towards the interior of the GUV, as indicated in the top image. (F) Sorting of ezrinTD in ABBA I-BAR domain-induced tubules at two different membrane coverages of ezrinTD. Measurements were collected from N = 19 GUVs, n = 3 experiments. Scale bars, 5 μm ϕ_v_: surface fraction of ezrinTD and ezrinWT on the GUVs. In (C, D and F), blue lines indicate *S* = 1. The following figure supplements are available for figure 6: **Figure supplement S9**. Sorting of IRSp53 I-BAR domain on self-induced membrane tubules and PIP_2_ enrichment in these tubules. **Figure supplement S10**. Colocalization of ezrinTD or ezrinWT with the I-BAR domain in tubules. **Figure supplement S11**. Direct interaction of ezrinTD with the I-BAR domain of ABBA.

It has been shown that BAR-domain proteins, including I-BAR, induce the formation of stable PIP_2_ clusters (Saarikangas et al., 2009) (Zhao et al., 2013) that can recruit downstream partners (Picas et al., 2014). It is thus conceivable that local PIP_2_ clusters induced by IRSp53 I-BAR domain facilitate ezrin enrichment. Indeed, we observed PIP_2_ enrichment in the I-BAR domain induced tubules, with a sorting ratio *S* up to 3 (Fig. S9C). As a control, we observed no sorting induced by the I-BAR domain for another fluorescent lipid GM1* (BODIPY-FL C5-ganglioside GM1) (Fig. S9C). Notably, this PIP_2_ enrichment is weaker than the enrichment of ezrinTD with *S* up to 8 (Fig. 6C) (given the tubule radii were set by the curvature of the I-BAR domain (Prevost et al., 2015), to compare PIP_2_ sorting and ezrin sorting at *ϕ_v_* < 2%, we performed *t*-test and obtained *p* = 1.2 × 10^−17^). This observation thus shows that the direct interaction between ezrin and the I-BAR domain enhances ezrin enrichment in I-BAR domain induced tubules.

MIM and ABBA, two other I-BAR domain proteins, were shown to colocalize with ezrin at the edge of transendothelial cell tunnels, which have a high negative curvature of about 1/30 nm (Stefani et al., 2017). Similarly to our measurements with the I-BAR domain of IRSp53, we found that ezrinTD interacts with ABBA I-BAR domain directly (Fig. S11) and ezrinTD was enriched in ABBA I-BAR domain-induced membrane tubules, with *S* > 1 for *ϕ_v_* < 2% (Fig. 6 E and F). Therefore, our data indicates that the enrichment of ezrinTD via ABBA I-BAR domain accounts for ezrinTD enrichment at transendothelial cell tunnels.

Taken together, our results evidence that ezrin enrichment in negatively curved membranes requires curvature-sensitive partners such as I-BAR domain proteins.

## Discussion

In cells, ezrin is associated with actin-rich membrane protrusions where the membranes have negative curvature, with intracellular vesicles where the membranes have positive curvature, and with the cell cortex where the membrane is flat. Using biomimetic model membranes having different curvatures combined with cell biology approaches, we report here the enrichment of ezrin on curved/tubular membranes via mechanisms involving the specific conformation of ezrin for positive curvature and the binding of ezrin to I-BAR domains for negative curvature.

### Ezrin forms anti-parallel assemblies that tether PIP_2_-containing membranes

It has been proposed that in solution, both ezrinWT and ezrinTD form anti-parallel homodimers via the association of the FERM domain of one ezrin monomer and of the C-ERMAD of the other monomer, bringing the α-helical coiled-coils of the two monomers together (Chambers and Bretscher, 2005) (Phang et al., 2016). We show that in the presence of membranes containing PIP_2_, both ezrinWT and ezrinTD self-assemble into densely packed, brush-like structures that tether two adjacent membranes. Upon PIP_2_ binding, the FERM domain of ezrin interacts with the membrane; the intermolecular interaction between the N- and C-ERMAD terminals of two opposite ezrin molecules drives the formation of anti-parallel assemblies, thereby inducing the “zipping” of two ezrin-coated membranes (Fig. 1). Our observations agree with previous reports indicating that PIP_2_ binding weakens the intramolecular interaction between the FERM and C-ERMAD of ezrinWT (Pelaseyed et al., 2017) (Shabardina et al., 2016). This reduction of the intramolecular interaction allows trans-intermolecular interactions between the FERM and C-ERMAD of two ezrinWT molecules on adjacent membranes, thus inducing zipping of the two adjacent membranes. Phosphomimetic mutation further reduces the intramolecular interaction (Shabardina et al., 2016) (Zhu et al., 2007), thus decreasing the intermolecular interaction, as evidenced with our dual micropipette aspiration experiments (Fig. 1F). We observed *in vitro*, for the first time, a tethering activity of ezrin not only at high protein concentration by Cryo-EM but also at low protein concentration by dual-GUV experiments, where ezrin density increases locally at the GUV-GUV interface upon zipping. This suggests that ezrin can cluster at the tethering interface even at low bulk concentration. Although in cells the concentration of ezrin is unknown, this activity may contribute to the juxtaposition of the membranes before fusion in physiological processes, such as the fusion of tubulovesicles with the apical canalicular membrane of parietal cells (Hanzel et al., 1991) in an ezrin-dependent manner to regulate acid secretion(Zhou et al., 2005) (Tamura et al., 2005), or the extension of the lumen of the excretory canal during development of C. elegans depending on an ezrin orthologue, ERM-1 (Khan et al., 2013).

### The conformation of ezrin phosphomimetic mutant accounts for its positive membrane curvature sensing

We find that on PIP_2_-containing membranes, ezrinWT and ezrinTD have distinct conformations. Consistent with previous AFM experiments on SLBs (Shabardina et al., 2016), the distance between the two FERM domains on tethered membranes is significantly reduced from 29 nm to 24 nm for the phosphomimetic mutant of ezrin. This difference in length of ezrinWT and ezrinTD essentially reflects a difference in the conformation of the molecules and thus possibly in their flexibility. Indeed, it was previously suggested that the α-helical region might exhibit different degrees of extensions depending on phosphorylation (Jayasundar et al., 2012) (Liu et al., 2007). The reduction of the GUV-tethering strength of ezrinTD as compared to ezrinWT indicates that a different conformation of the C-ERMAD of ezrin TD reduces its interaction with the N-ERMAD, and consequently the intermolecular interaction. We observe that ezrinTD, but not ezrinWT, recruits F-actin to PIP_2_-containing membranes by forming brush-like structures at the membranes. This observation confirms that although ezrinWT is partially open upon PIP_2_ binding (Pelaseyed et al., 2017) (Shabardina et al., 2016), its C-terminal actin-binding site remains inaccessible for F-actin binding, in agreement with previous reports (Fievet et al., 2004) (Fritzsche et al., 2014) (Zhu et al., 2007) (Gautreau et al., 2000). Finally, we observed that ezrinTD better conforms to the tubular membranes than ezrinWT and is moderately enriched onto positively curved membrane tubes *in vitro*, while ezrinWT is typically excluded. Late endosomes have a typical radius ranging between 100 to 200 nm, thus a curvature of about 2/200nm to 2/100nm (Huotari and Helenius, 2011). A clear difference was observed *in vitro* between ezrinTD and ezrinWT in this curvature range (Fig. 3 D and E). Although we cannot exclude that ezrin is recruited by a specific partner on endosomes such as the HOPS (HOmotypic fusion and Protein Sorting) complex (Chirivino et al., 2011), the positive curvature sensing of the ezrin phosphomimetic mutant observed here might facilitate phosphorylated ezrin association with positively curved endosomal membranes.

### Negative membrane curvature enrichment of ezrin requires its binding to I-BAR domain proteins

Ezrin is enriched in actin-rich membrane protrusions wherein the membranes have negative curvature. It has been proposed that PIP_2_ binding and the specific location of the kinase LOK that phosphorylates ezrin regulate the membrane localization of ezrin (Fievet et al., 2004) (Viswanatha et al., 2012). Here we show that ezrinWT and ezrinTD do not sense negative membrane curvature and that the interaction with I-BAR domain proteins is required to facilitate the enrichment of ezrinWT or ezrinTD on membrane protrusions. We further anticipate that the direct ezrin-I-BAR interaction together with the PIP_2_ clusters induced by the I-BAR domains (Saarikangas et al., 2009) (Zhao et al., 2013) synergistically enrich ezrin in the I-BAR domain-induced tubules. Given that IRSp53 contributes to the initiation of filopodia (Disanza et al., 2013), we propose that IRSp53 recruits ezrin to strengthen the binding of actin filaments to the plasma membrane that in turn facilitates filopodia growth.

In conclusion, our data corroborates the role of ezrin to maintain the mechanical cohesion of membranes with F-actin, thus facilitating actin-related cellular morphological changes. Furthermore, our work reveals new mechanisms underlying the recruitment of ezrin on different cellular membrane curvatures: the phosphomimetic mutation T567D of ezrin induces a more flexible conformation of ezrin that facilitates its interaction with positively curved membranes of some intracellular vesicles. In contrast, ezrin hijacks I-BAR domain proteins to accumulate on negatively curved membranes such as membrane protrusions and the edge of transendothelial cells’ tunnels where ezrin is known to play a major architectural function. Interestingly, another ezrin binding partner ERM Binding Protein 50 (EBP50) was shown to interact directly with IRSp53 (Garbett et al., 2013) thus could in turn recruit ezrin to microvilli. Considering that ezrin is one of the most abundant proteins in cells and given its actin-membrane linking function, it is presumably crucial for cells to have membrane-actin linkers that are by themselves curvature-insensitive to avoid inducing massive protrusions at the plasma membrane (Saarikangas et al., 2009).

## Materials and Methods

### Reagents

Total brain lipid extract (TBX, 131101P), brain l-α-phosphatidylinositol-4,5-bisphosphate (PIP_2_, 840046P), 1,2-distearoyl-sn-glycero-3-phosphoethanolamine-N-[biotinyl(polyethyleneglycol)-2000] (DSPE-PEG(2000)-biotin, 880129P), 1-oleoyl-2-{6-[4-(dipyrrometheneboron difluoride)butanoyl]amino},hexanoyl-sn-glycero-3-phosphoinositol-4,5-bisphosphate (TopFluor PIP_2_, 810184P), L-α-phosphatidylcholine (Egg, Chicken) (EPC, 840051), and galactocerebrosides were purchased from Avanti Polar Lipids/Interchim. BODIPY-TR-C5-ceramide, (BODIPY TR ceramide, D7540), BODIPY-FL C5-ganglioside GM1 (GM1*, B13950), BODIPYFL C5-hexadecanoyl phosphatidylcholine (HPC*, D3803), Alexa Fluor 488 C5-Maleimide (AX488), Alexa Fluor 546 C5-Maleimide (AX546), Alexa Fluor 633 C5-Maleimide (AX633), were purchased from Invitrogen. GloPIPs BODIPY^®^ TMR-PtdIns(4,5)P_2_, C16 (TMR PIP_2_, C45M16a) was purchased from Echelon. Non-muscle actin (Actin protein, >99% pure, human platelet, APHL99) was purchased from Cytoskeleton. Streptavidin-coated polystyrene beads (SVP-30-5) were purchased from Spherotech. TRITON X-100 α-[4-(1,1,3,3-Tetramethylbutyl)phenyl]-w-hydroxy-poly(oxy-1,2-ethanediyl) (Triton-anapoe, Anapoe-X-100, anatrace), mPEG-silane MW 2000 (mPEG-silan-2000) was purchased from Laysan Bio. Alexa Fluor 488 tagged phalloidin (AX488 phalloidin) was purchased from Interchim. β-casein from bovine milk (>98% pure, C6905) and other reagents were purchased from Sigma-Aldrich. For detecting endogenous ezrin in LLC-PK1 cells, anti-ezrin antibodies from M. Arpin laboratory was used (Algrain et al., 1993). For immunoprecipitation: anti-ezrin antibodies (cat# 610602, BD transduction laboratories) and anti-GFP antibodies (cat#11814460001, Roche). HRP coupled anti-mouse antibodies are from The Jackson laboratory and Cy3 coupled anti-rabbit antibodies are from The Jackson laboratory.

### Plasmid construction

cDNA coding for either full-length human ezrin wild-type (ezrinWT) or its phosphomimetic version, ezrin T567D (ezrinTD) (Gautreau et al., 2000) were cloned into a pENTR vector (Invitrogen) by BxP recombination. pET28-N-His-SUMO plasmids (N-terminal His tag and SUMO fusion vector) containing the cDNA coding for either ezrinWT or ezrinTD were constructed by Gibbson Assembly (New England BioLabs) and used for the production of recombinant proteins. To ensure fluorescent maleimide labeling of recombinant ezrinWT and ezrinTD two extra cysteine residues were inserted at the C-terminal using quick-change site-directed mutagenesis (Agilent Technologies), based on the previous report (Blin et al., 2008). All the constructions were finally verified by sequencing.

### Protein purification and labeling

Purifications of the full-length ezrinWT, ezrinTD and the FERM domain of ezrin were performed based on a previously described procedure (Braunger et al., 2013). His-tagged proteins were expressed in *E. coli* B121 codon plus (DE3)-RIL cells induced by 0.5 mM Isopropyl β-D-1-thiogalactopyranoside (IPTG) for 20 hr at 20°*C*. In the following, all steps were performed at 4°*C* or on ice. Bacterial pellets were collected by centrifugation at 5000g (JLA 9.1000 rotor) for 20 min, resuspended in lysis buffer (40 mM HEPES pH 7.2, 300 mM NaCl, 5 mM β-Mercaptoethanol, one protease inhibitor tablet (complete ULTRA tablets EDTA free) and 1mg.mL^−1^ lysozyme), incubated for 30 min at 4°C and tip sonicated on ice for 4 min at 10sec/10sec, power 35%. Clear lysate was obtained by centrifugation of the lysate at 48000g (JA 25.50 rotor) for 30min, followed by filtered with filtration unit Stericup (0.22μm). Purified proteins were isolated by chromatography using 2xlml Protino^®^ Ni-NTA Columns. The Ni-NTA column was equilibrated in equilibration buffer (40 mM HEPES pH 7.2, 300 mM NaCl, 5 mM β-Mercaptoethanol, 10 mM Imidazole) and bound proteins were eluted in elution buffer (40mM HEPES pH 7.2, 300 mM NaCl, 5 mM β-Mercaptoethanol, 300 mM Imidazole) with a linear gradient from 10% to 100% of the elution buffer. To cleave the His tag, the eluted proteins were dialyzed overnight in dialysis buffer (40 mM HEPES pH 7.2, 300 mM NaCl, 5 mM β-Mercaptoethanol) containing SUMO protease. Dialyzed proteins were then purified with a Ni-NTA column equilibrated in the dialysis buffer. Finally, the eluted proteins were buffer exchanged into the labeling buffer (20 mM HEPES pH 7.2, 50 mM KC1, 0.1 mM EDTA) on a PD10 column (GE Healthcare). If not continuing the following labeling steps, pure proteins were supplemented with 2 mM β-Mercaptoethanol and 0.1% methylcellulose, snapped frozen in liquid nitrogen and stored at −80°*C*.

EzrinTD, ezrinWT and the FERM domain of ezrin labeling were performed right after the last elution step. A 5 molar excess of Alexa Fluor maleimide dyes was added to the pure proteins and allow to label at 4°*C* overnight. The labeling was quenched by supplementing 2 mM β-Mercaptoethanol in the reaction and the dye removed on a PD10 column (GE Healthcare) into the storage buffer (20 mM Tris pH 7.4, 50 mM KC1, 0.1 mM EDTA, 2 mM β-Mercaptoethanol). Finally, Alexa Fluor labeled pure proteins were supplemented with 0.1% methylcellulose, snapped frozen in liquid nitrogen and stored at −80°*C*.

Muscle actin was purified from rabbit muscle and isolated in monomeric form in G-buffer (5 mM Tris-Cl^−^, pH 7.8, 0.1 mM CaCl_2_, 0.2 mM ATP, 1 mM DTT, 0.01% NaN_3_) as previously described (Spudich and Watt, 1971). Recombinant mouse IRSp53 I-BAR domain and ABBA I-BAR domain were purified and labeled as previously described (Prevost et al., 2015) (Saarikangas et al., 2009). Gelsolin was purified as previously described (Le Clainche and Carlier, 2004).

### Flow cytometry experiment

GUVs containing BODIPY TR ceramide were incubated with AX488 ezrin for at least 20 min at room temperature before flow cytometry measurement (BD LSRFortessaTM cell analyzer, BD Bioscience). Data was collected by BD FACSDivaTM software (BD Bioscience) and analyzed by using Flowing Software 2.5.1 (www.flowingsoftware.com). Fig. S1F shows an exemplifying figure for gating

### Measuring membrane surface fraction of proteins

We measured protein surface density on GUV membranes (number of proteins per unit area) by performing a previously established procedure (Sorre et al., 2012) (Sorre et al., 2009). We related the fluorescence intensity of the fluorescent dye used to label proteins (AX488) to that of a fluorescent lipid (BODIPY FL-C5-HPC, named HPC*). We can measure fluorescence intensity of HPC* on GUV membranes at a given HPC* membrane fraction. Thus, the surface density of the protein on membranes is *n_protein_* = 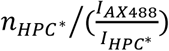, where 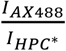 is the factor accounting for the fluorescence intensity difference between HPC* and AX488 at the same bulk concentration under identical image acquisition condition. The area density of HPC*, *Φ_HPC*_*, can be related to its fluorescence intensity, 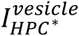, by measuring fluorescence intensities of GUVs composed of DOPC supplemented with different molar ratios of HPC* (0.04-0.16 mole%) and assuming lipid area per lipid is 0.7 nm^2^ (1120 – 4480 HPC* per μm^2^). As such, 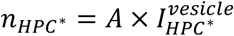, where *A* is a constant depending on the illumination setting in the microscope. We then obtained the surface density of the protein as 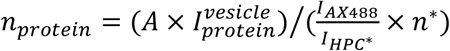, where *n*\* is the degree of labeling for the protein of interest. Finally, we obtained the surface fraction of the protein *Φ_protein_* = *n_protein_* × *a_protein_*, where *a_protein_* is the area of a single protein on membranes. *a_ezrin_* ≅ 20 nm^2^ was obtained by EM analysis as shown in Figure 1C, and *a_I–BAR domain_* ≅ 50 nm^2^ (Millard et al., 2005).

To obtain protein/membrane fluorescence intensity in tube pulling experiments, we manually defined the region of interest, a rectangle, around the membrane of a GUV or around the membrane nanotube of the GUV such that the membrane/tube was horizontally located at the center of the rectangle. We then obtained an intensity profile along the vertical direction of the rectangle by calculating the mean fluorescence intensity of each horizontal line of the rectangle. To account for protein fluorescence singles in the inner GUV when encapsulating or in the outer GUV when injecting, the background protein intensity was obtained by calculating the average value of the mean of the first 15 intensity values from the top of the rectangle intensity profile and the mean of the first 15 intensity values from the bottom of the rectangle intensity profile. The background membrane intensity was obtained by calculating the mean of the first 15 intensity values from the top of the rectangle intensity profile. Finally, the protein/membrane fluorescence intensity was obtained by subtracting the background intensity from the maximum intensity value in the intensity profile.

### F-actin preparation

Muscle F-actin was pre-polymerized for at least 1hr at RT from Mg-ATP-actin in the presence of gelsolin at a gelsolin:actin ratio of 1:1380 to obtain F-actin with controlled lengths. As such, the resulting muscle F-actin has lengths of a few μm (Harris and Weeds, 1984). The same procedure was performed for non-muscle F-actin, except that the gelsolin:actin ratio is 1:2069. To visualize actin, both muscle and non-muscle F-actin was labeled with equal molar amount of AX488 phalloidin.

### Glass passivation

For all experiments, micropipettes for holding GUVs and microscope slides and coverslips were washed with water and ethanol followed by passivated with a □-casein solution at a concentration of 5 g.L^−1^ for at least 5 min at RT. For ezrin/I-BAR domain GUV experiments, microscope slides and coverslips were cleaned by sonication with water, ethanol and acetone for 10 min, 1M KOH for 20 min, and then water for 10 min. Observation chambers were assembled and passivated with □-casein solution (5 g.L^−1^ in PBS buffer) and then with mPEG-silan-2000 solution (5mM in DMSO), each for at least 5 min.

### Cell culture and transfection

LLC-PK1 cell line was obtained from ATCC (CCL 101; American Type Culture Collection, Rockville, MD) (Coscoy et al., 2002) and HeLa cell line was from B. Goud laboratory (Echard et al., 1998). LLC-PK1 cell line authentication was performed by M. Arpin laboratory (Coscoy et al., 2002) and HeLa cell was not authenticated. Cell lines were tested negative for mycoplasma contamination. LLC-PK1 cells and HeLa cells were cultivated at 5% CO_2_ and at 37°*C* in DMEM supplemented with 10% (v/v) serum and 5% (v/v) penicillin/stretomycin. Transient transfection of LLC-PK1 and HeLa cells were performed with X-tremeGENE HP and X-tremeGENE 9 (Sigma-Aldrich), respectively, according to the manufacturer’s instruction. Co-immunoprecipitation and mass spectrometry analysis of GFP-tagged proteins was performed by using GFP-Trap^®^ (Chromotek) according to the manufacturer’s instruction.

### GUV lipid compositions and buffers

In the following experimental procedures, we use “ezrin” to refer to ezrinTD and ezrinWT. Lipid compositions for GUVs were total brain lipid extract (TBX) (Yu et al., 2006) supplemented with 5 mole% brain PIP_2_, 0.025-0.5 mole% DSPE-PEG(2000)-biotin and 0.5-1 mole% BODIPY TR ceramide for ezrin tube pulling experiments. TBX supplemented with 0.1 mole% DSPE-PEG(2000)-biotin and 0.5 mole% BODIPY TR ceramide with and without 5 mole% brain PIP_2_ for testing ezrin-PIP_2_ binding. TBX supplemented with 5 mole% brain PIP_2_, 0.1 mole% DSPE-PEG(2000)-biotin and 0.5 mole% BODIPY TR ceramide for assessing ezrin-membrane binding affinity by using confocal microscopy. TBX supplemented with 5 mole% brain PIP_2_, and 0.5 mole% BODIPY TR ceramide for assessing ezrin-membrane binding affinity by using flow cytometry. TBX supplemented with 5 mole% brain PIP_2_, and 0.5 mole% BODIPY TR ceramide for GUV-ezrin tethering assay. TBX supplemented with 5 mole% brain PIP_2_, 0.2 mole% DSPE-PEG(2000)-biotin and 0.5 mole% BODIPY TR ceramide for biotin-streptavidin tethering assay. TBX supplemented with 5 mole% brain PIP_2_, and 0.5 mole% BODIPY TR ceramide for AX488 labeled ezrin and unlabeled I-BAR domain recruitment experiments. TBX supplemented with 5 mole% brain PIP_2_, and 0.8 mole% Rhodamine-PE ceramide for AX633 labeled ezrin and AX488 labeled I-BAR domain recruitment experiments. TBX supplemented with 4.5 mole% brain PIP_2_, 0.5 mole% TopFluor PIP_2_, 0.2 mole% DSPE-PEG(2000)-biotin and 0.5 mole% BODIPY TR ceramide for TopFluor PIP_2_ tube pulling experiments. TBX supplemented with 4.8 mole% brain PIP_2_, 0.2 mole% TMR PIP_2_, 0.2 mole% DSPE-PEG(2000)-biotin and 0.8 mole% GM1* for TMR PIP_2_ tube pulling experiments. TBX supplemented with 4.5 mole% brain PIP_2_, 0.5 mole% TopFluor PIP_2_, and 0.5 mole% BODIPY TR ceramide, TBX supplemented with 4.2 mole% brain PIP_2_, 0.8 mole% TopFluor PIP_2_, and 0.5 mole% BODIPY TR ceramide for PIP_2_-I-BAR domain induced sorting experiments. TBX supplemented with 5 mole% brain PIP_2_, 0.8 mole% GM1*, and 0.5 mole% BODIPY TR ceramide for GM1*-I-BAR domain induced sorting experiments. DOPC supplemented with 10mole% DGS-NTA(Ni) for preparing Ni-GUVs.

The salt buffer outside GUVs was 60 mM NaCl and 20 mM Tris pH 7.5, except for ezrin encapsulation experiments where the buffer was 300 mM NaCl and 20 mM Tris pH 7.5 to detach ezrin from binding on the outer leaflet of GUVs, and for the dilution experiments where GUVs containing ezrin was diluted in 60 mM NaCl, 430 mM glucose and 20 mM Tris pH 7.5 this buffer. The salt buffer inside GUVs was 50 mM NaCl, 20 mM sucrose and 20 mM Tris pH 7.5, except for GUVs encapsulating ezrin where the buffer was 60 mM NaCl, 430 mM sucrose and 20 mM Tris pH 7.5, and for Ni-GUV experiments where 157 mM sucrose solution was used.

### GUV preparation

For all experiments, GUVs were prepared by electroformation on platinum electrodes under a voltage of 0.25 V and a frequency of 500 Hz overnight at 4°C in a physiologically relevant salt buffer (Méléard et al., 2009), except for flow cytometry experiments and for preparing Ni-GUVs. For flow cytometry experiments, GUVs was prepared by using polyvinyl alcohol (PVA) gel-assisted method in a salt buffer (60 mM NaCl) at room temperature for 1 hour as described previously (Weinberger et al., 2013). Ni-GUVs were prepared by using electroformation on ITO-coated plates under a voltage of 1 V and a frequency of 10 Hz for 1 hour at room temperature in a sucrose buffer (154 mM sucrose) as described previously (Montes et al., 2007).

For GUVs encapsulating ezrin, 0.2-0.5 μM of ezrin was present during GUV growth. As such, the resulting GUVs have ezrin binding on both the inner and outer leaflets of the GUV membranes. Since ezrin binds to PIP_2_ containing membranes via electrostatic interactions, screening these interactions should result in ezrin desorbing from the membranes. Indeed, when placing GUVs coated with ezrin in a high ionic strength buffer (300 mM NaCl) while keeping the osmotic pressure of the GUVs balanced, a nearly complete depletion of ezrin form GUVs was observed (Fig. S8).

### Tube pulling experiments

Tube pulling experiments were performed in a setup comprised a Nikon C1 confocal microscope equipped with a X60 water immersion objective, micromanipulators for positioning micropipettes and optical tweezers as described previously (Sorre et al., 2009). To pull a tube, a GUV was held by a micropipette, brought into contact with a streptavidin-coated bead trapped by the optical tweezers, and then moved away from the bead. The tube radius *R* was measured by using the ratio of lipid fluorescence intensity on the tube and on the GUV as 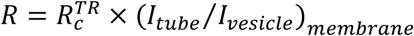, where *R_c_* = 200 ± 50 nm is the previously obtained calibration factor for using BODIPY TR ceramide as lipid fluorescence reporter in the same setup by performing a liner fit of membrane fluorescence ratio (*I_tube_/I_vesicle_*)*_membrane_* and lipid radii *R* measured by *R* = *f*/(4*πσ*), where *f* are forces applied by the optical tweezers to sustain the tubes and *σ* are the membrane tensions controlled by the micropipette hold the GUVs, as previous reports (Sorre et al., 2009) (Prevost et al., 2015). For experiments where GM1* lipids were used as lipid reporter, we obtained the calibration factor 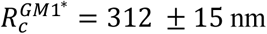.

### GUV-tethering experiments

For ezrin tethering, GUVs were incubated with 0.4-3 μM of AX488 ezrin in solution. For biotin-streptavidin tethering, GUVs incorporating with 0.2 mole% biotinylated lipids were incubated with 0.4 μM or 0.8 μM of AX488 streptavidin in solution. In a typical experiment, two GUVs coated with proteins were held by micropipettes and brought into contact. We then decreased stepwise the membrane tension of the test GUV. The contact size of the two micropipette held GUVs is determined by the force balance at the contact zone: σ cos(θ) = σ − γ, where θ is the contact angle of the two GUVs, σ is the membrane tension of the test GUV and γ is the tethering energy at the contacting zone (see Fig. S3C for the schematic) (Franke et al., 2006). The number of proteins at the tethering zone is estimated by using fluorescence signals of the proteins, assuming all ezrin at the tethering zone contributes to the membrane tethering. This is a reasonable assumption, given that in our EM observation, we observed densely packed ezrin oligomers in-between bilayers. To obtain binding energy per ezrin bond, we assumed that ezrin tethers the membranes in its dimer form, and thus divided the tethering energy by the number of ezrin dimers at the tethering zone. The binding energy per biotin-streptavidin bond was obtained by dividing the tethering energy by the number of streptavidin at the tethering zone. Samples were observed by a X60 water immersion objective with an inverted confocal microscope (Nikon TE2000 microscope equipped with eC1 confocal system).

### Ezrin/I-BAR domain GUV assay and its fluorescence intensity quantification

For IRSp53 I-BAR domain-GUV experiments, GUVs were incubated with I-BAR domain at a concentration of 0.02-0.5 μM (containing 30 mole% of AX488 IRSp53 I-BAR domain) for at least 30 min at RT before observation. For ezrin/I-BAR domain GUV experiments, GUVs were first incubated with ezrin (0.05-2 μM) for at least 15 min at RT, and then I-BAR domain (0.05-2 μM) was added into the ezrin-GUV mixture. Samples were observed by a X100 oil immersion objective with an inverted spinning disk confocal microscope (Nikon eclipse Ti-E equipped with a EMCCD camera, QuantEM, Photometrics).

Here, to obtain protein and membrane fluorescence intensities on a vesicle membrane or on the corresponding membrane tube to calculate (*I_tube_/I_vesicle_*)_*membrane*_ and (*J_tube_/I_vesicle_*) _*protein*_, we manually defined the region of interest, a line with a width of 6 pixels drawn perpendicularly cross the membrane region of interest. We then obtained the intensity profile of the line where the x-axis of the profile is the length of the line and the y-axis is the averaged pixel intensity along the width of the line. The background intensity was obtained by calculating the mean value of the sum of the first 10 intensity values and the last 10 intensity values of the intensity profile. Finally, the protein and membrane fluorescence intensities were obtained by subtracting the background intensity from the maximum intensity value in the intensity profile. This image process was performed by using Fiji (Schindelin et al., 2012).

To ensure we measured the protein and membrane fluorescence intensities on tubes that are in focus, we typically recorded 60 images of a GUV with 100 millisecond exposure time and at a frame interval of around 0.5 second and manually selected tubes that are in focus.

### Line profile along cellular protrusions

A line with a width of 4 pixels were manually drawn along the protrusions and the corresponding intensity profiles were obtained by using Fiji (Schindelin et al., 2012).

### Cryo-electron microscopy

LUVs were prepared by detergent elimination (Rigaud et al., 1998). Briefly, a lipid mixture of TBX supplemented with 5 mole% PIP_2_ was dried with argon gas and placed under vacuum for at least 3hrs. The dried lipid film was resuspended in a salt buffer (60 mM NaCl and 20 mM Tris pH 7.5) at a concentration of 1 g.L^−1^. 20 mL of Triton-anapoe (10% w/v) was added into the lipid suspension and was gradually eliminated throughout the addition of biobeads in a stepwise manner to generate LUVs (20 mg of biobeads were added for an overnight incubation at 4°*C* followed by two subsequent additions of biobeads, 20 mg and 40 mg, the next morning).

LUVs were incubated with ezrin and/or F-actin at varying concentrations for at least 15 minutes at RT before vitrification. The samples were vitrified on copper holey lacey grids (Ted Pella) using an automated device (EMGP, Leica) by blotting the excess sample on the opposite side from the droplet of sample for 4 seconds in a humid environment (90 % humidity). Imaging was performed on a LaB6 microscope running at 200 kV (Technai G2, FEI) and equipped with a 4KX 4K CMOS camera (F416, TVIPS). Automated data collection for 2D imaging as well as tilted series collection for cryo-tomography were carried out with the EMTools software suite.

Galactocerebroside nanotubes were prepared following the protocol given by (Dang et al., 2005). Briefly, galactocerebroside (Galact (β) C24 :1 Cer) was mixed with 5 mole% PIP_2_ and 15 mole% EPC in chloroform and methanol at 10 g.L^−1^. After extensive drying, the lipid film was re-suspended in 20mM HEPES pH7 and imidazole 200mM at 5 g.L^−1^. The solution was stirred at room temperature for one hour.

### Two dimensional image processing for EM images

918 and 321 square boxes of 253 pixels were hand-picked from the TD ezrin and WT ezrin images respectively using the boxer tool from the EMAN software suite (Ludtke et al., 1999). Subsequent processing was carried out using SPIDER (Frank et al., 1996). After normalization of the particles, a non-biased reference-free algorithm was used to generate 10 classes. 1124 boxes of 99 pixels were selected to average the CERMAD globular domain of ezrin and the same protocol described above was carried out. To enhance the signal-to-noise ratio, we performed two dimensional single particle analysis by selecting pieces of stacks (square boxes) comprising both bilayers and the protein material in between (900 and 320 boxes for ezrinTD and ezrinWT, respectively). Class averages were then generated by statistical analysis algorithms. The class averages were then used individually to measure the distance between globular FERM domains within tethers. The resulting value is the average from the distances measured for each class and the error is given by the standard deviation.

### Cryo-electron tomography: data collection and image processing

Cryo-tomography was performed using a LaB6 microscope running at 200 kV (Technai G2, FEI) and equipped with a 4KX 4K CMOS camera (F416, TVIPS). Prior to the vitrification of the sample 10 nm gold beads were added in solution to be subsequently used as fiducials. Data collection was carried out using the EMTool (TVIPS) software suite. Tilted series were acquired from −60 to 60 degrees using a saxton angular data collection scheme. Individual images were collected with a 0.8 electrons per Å^2^ for a total dose of less than 70 electrons per Å^2^. Imod was primarily used for data processing and alignment for individual images. The reconstructions were performed using either IMOD (Weighted back projection) or Tomo3D (SIRT). The segmentation of the volumes was performed manually using IMOD.

### Immunofluorescence and immunoprecipitation

For immunofluorescence labeling, transfected LLC-PK1 cells were fixed in 3% paraformaldehyde in phosphate-buffered saline supplemented with 1 mM MgCl_2_ and 1mM CaCl_2_ (PBS^+^) for 20 min, washed with PBS^+^ and then incubated with 0.5% Triton X100 in PBS^+^ incubated with rabbit anti-ezrin antibodies for 1 hr, washed with PBS^+^, incubated with secondary fluorescent antibodies (Cy3) for 1 hr, and washed with PBS^+^.

For immunoprecipitation, HeLa cells transfected with plasmids encoding GFP, GFP-I-BAR domain or GFP-IRSp53 were washed in PBS, trypsinized, and pelleted by centrifugation at 4°*C*. Pelleted cells were then lysed in lysis buffer (25 mM Tris pH 7.5, 50 mM NaCl, 0.1% NP40, and protease inhibitor mix). Lysates was passed 3 times through a 25g syringe, incubated on ice for 1 hr to extract membrane bound proteins and then centrifuged for 10 min at 10,000 rpm to remove insoluble material and nucleus. The clean lysates were then incubated with GFP-Trap beads at 4°*C* under rotation for 3 hr. The lysate-bead mixtures were washed 2 times in lysis buffer after centrifugation at 1850 rpm for 2 min at 4°*C*. The remaining buffer after the last centrifugation was removed by using a syringe. The dry lysate-bead pellet was resuspended in Laemmli buffer and processed for Western blotting following standard protocols.

### Statistics

All notched boxes show the median (central line), the 25th and 75th percentiles (the edges of the box), the most extreme data points the algorithm considers to be not outliers (the whiskers), and the outliers (circles).

## Author Contributions

P. B., E. C. and F.-C. T. designed research. E. L. and P. L. provided conceptual advice. F.-C. T. performed *in vitro* and cell biology experiments, A.B. performed EM experiments, S. M.-L. performed mass spectrometry experiments, M.-C. T. performed experiments of transendothelial cell tunnels. F.-C. T. and A.B. analysed data. H. B. purified ezrin, and J. M. purified I-BAR domains. L. P. provided ezrin constructs and Y. S. provided GFP-IRSp53 plasmid. F.-C. T., A. B., E. L., E. C and P. B. wrote the manuscript. All authors discussed the results and provided feedback on the manuscript.

## Acknowledgments

We thank B. Goud for insightful discussions, C. Le Clainche (Institute for Integrative Biology of the Cell, Gif-sur-Yvette, France) and J. Pernier for providing actin and advice on actin reconstitution, F. Brochard for helping on analysis of the GUV-tethering assay, F. Di Federico for handling plasmids, C. Prévost for her help for the optical tweezers setup and data analysis, A. Di Cicco and D. Levy for EM image acquisition and galcer tube preparation, N. de Franceschi for helping on the FACS experiments, M. Henderson for carefully reading of the manuscript. The authors greatly acknowledge the Cell and Tissue Imaging (PICT-IBiSA), Institut Curie, member of the French National Research Infrastructure France-BioImaging (ANR10-INBS-04). This work was supported by Institut Curie, Centre National de la Recherche Scientifique (CNRS), the Agence Nationale pour la Recherche (grant ANR-15-CE18-0016-03 to P.B.), the European Research Council (E.C. and P.B. partners of the advanced grant, project 339847) and the Human Frontier Science Program Organization (RGP0005/2016 to P.B.). E. C. and P. B.s groups belong to the CNRS consortium CellTiss, to the Labex CelTisPhyBio (ANR-11-LABX0038) and to Paris Sciences et Lettres (ANR-10-IDEX-0001-02). F.-C. Tsai was funded by the EMBO Long-Term fellowship (ALTF 1527-2014) and Marie Curie actions (H2020-MSCA-IF-2014, project membrane-ezrin-actin).

---

**Figure SI.**
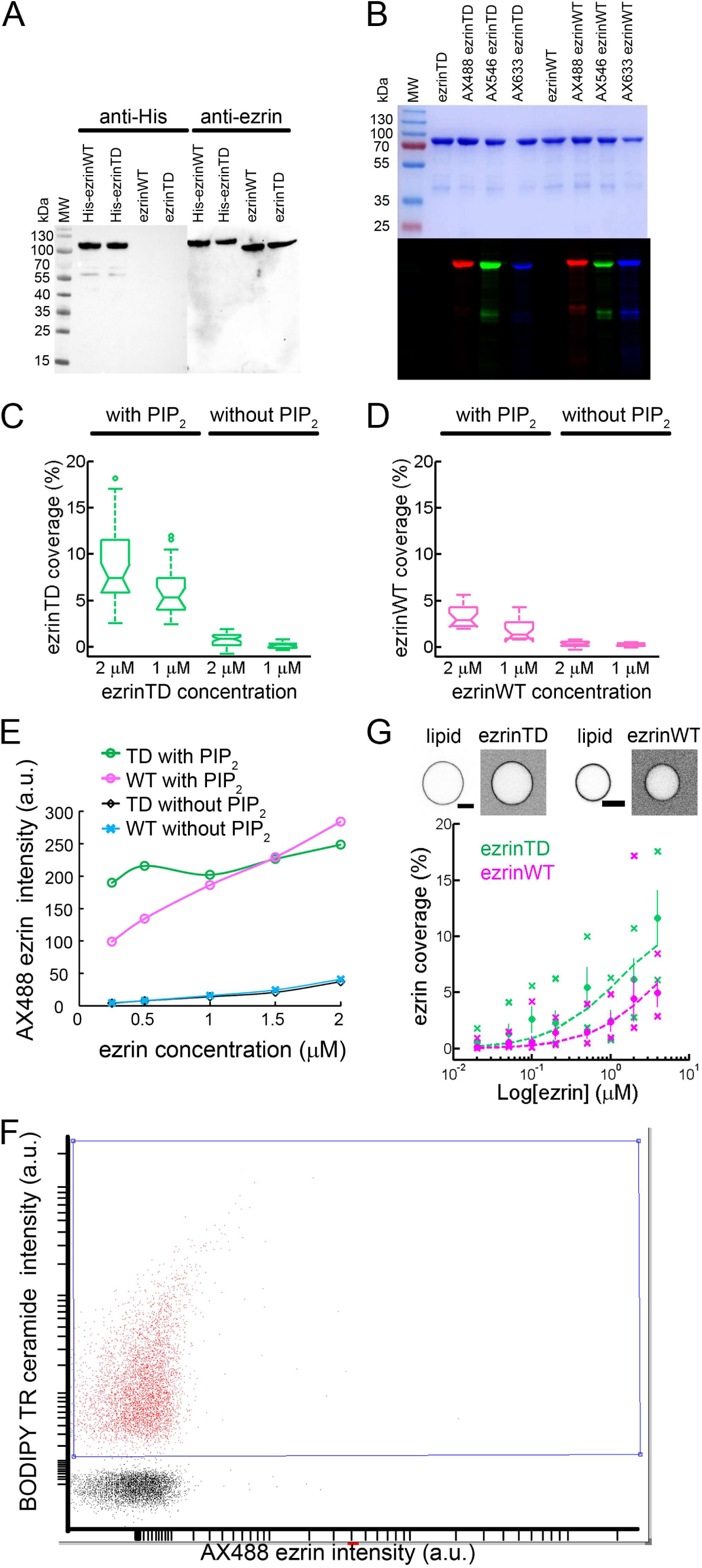
Analysis of purified recombinant ezrinTD and ezrinWT, and their binding to PIP_2_-containing membranes, related to Figure 1. (A) Western blots of ezrinTD and ezrinWT before and after SUMO proteolysis of the His_6_ tag analyzed with anti-His or anti-ezrin antibodies. MW: molecular weight protein markers. (B) SDS-PAGE analysis of Alexa dye-labelled ezrinTD and ezrinWT detected by Coomassie blue staining (top) and fluorescence illumination (bottom). (C and D) GUV surface fraction covered with ezrinTD (C) and ezrinWT (D) in the presence and absence of PIP_2_. N = 34, **22**, 39, 30 and N = 18, 15, 24, 20 GUVs for GUVs with PIP_2_ at 2 μM and 1 μM protein concentrations and GUVs without PIP_2_ at 2 μM and 1 μM protein concentrations, for ezrinTD and ezrinWT, respectively. n = 1 experiment. Measurements were taken from distinct samples. (E) Flow cytometry analysis of ezrinTD and ezrinWT binding to GUVs with and without PIP_2_. Number of events for all conditions N > 7600. (F) An example of gating on GUVs analyzed by flow cytometry. 100% events is 10000. (G) (Top) Representative confocal images of ezrinTD and ezrinWT on GUVs containing 5 mole% PIP_2_. Scale bars, 5 μm. Inverted grayscale images were shown. (Bottom) Representative measurement of the surface fraction of ezrinTD and ezrinWT at various ezrin bulk concentrations for n = 2 experiments. Statistics shown here: N = 43-78 and N = 69-79 GUVs per condition for ezrinTD and ezrinWT, respectively, for one of the two experiments. Measurements were taken from distinct samples. Error bars indicate standard deviations, circles are mean and X symbols are maximum and minimum for each condition. Dash lines are data fitting curves.

**Figure S2.**
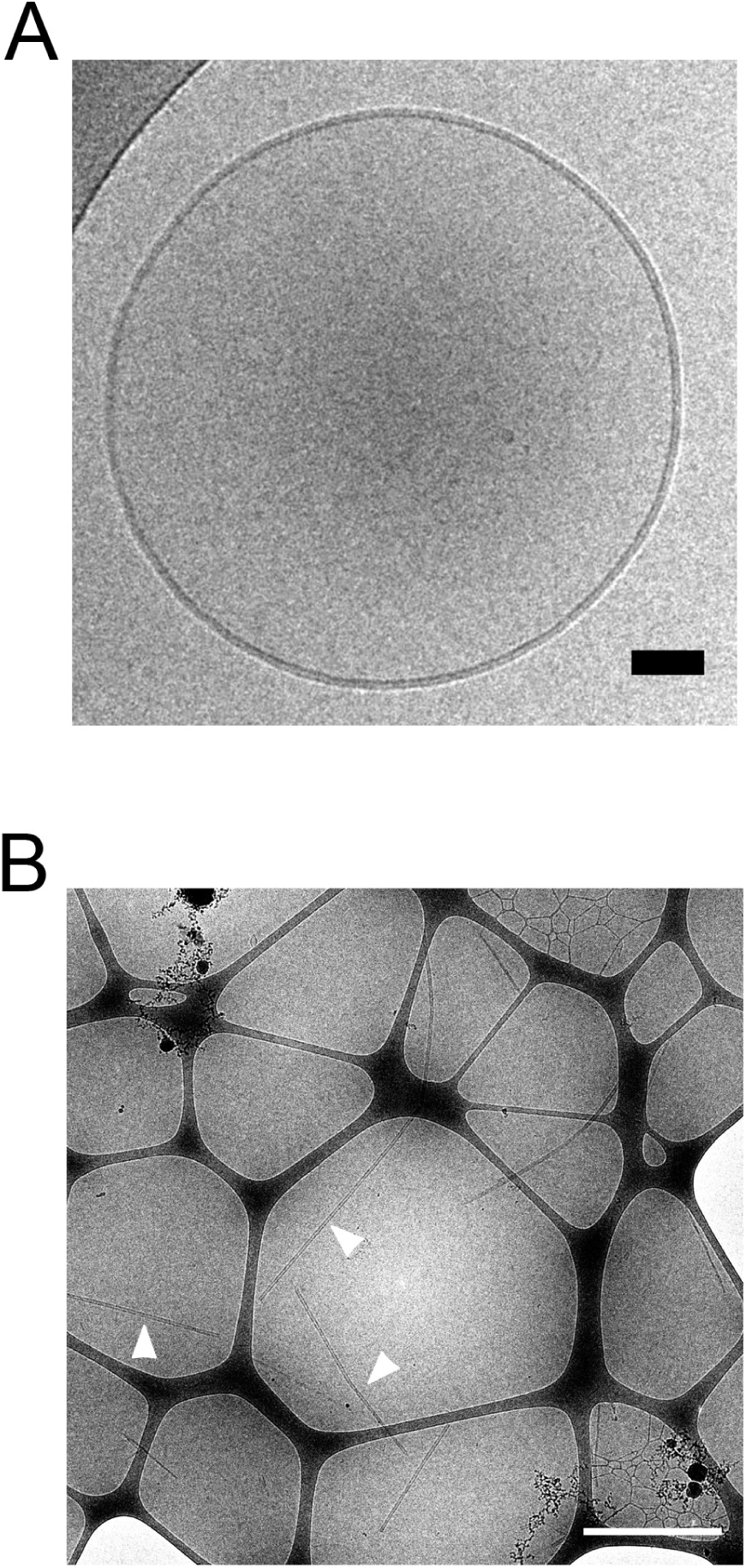
Representative cryo-electron micrograph of LUVs and galactocerebroside nanotubes in the absence of ezrin, related to Figure 1 and Figure 3. (A) LUV in the absence of ezrin. Scale bar, 50 nm. (B) Galactocerebroside nanotubes in the absence of ezrin. Arrowheads indicate some of the nanotubes. Nanotube concentration: 0.1 g.L^−1^. Scale bar, 1 μm.

**Figure S3.**
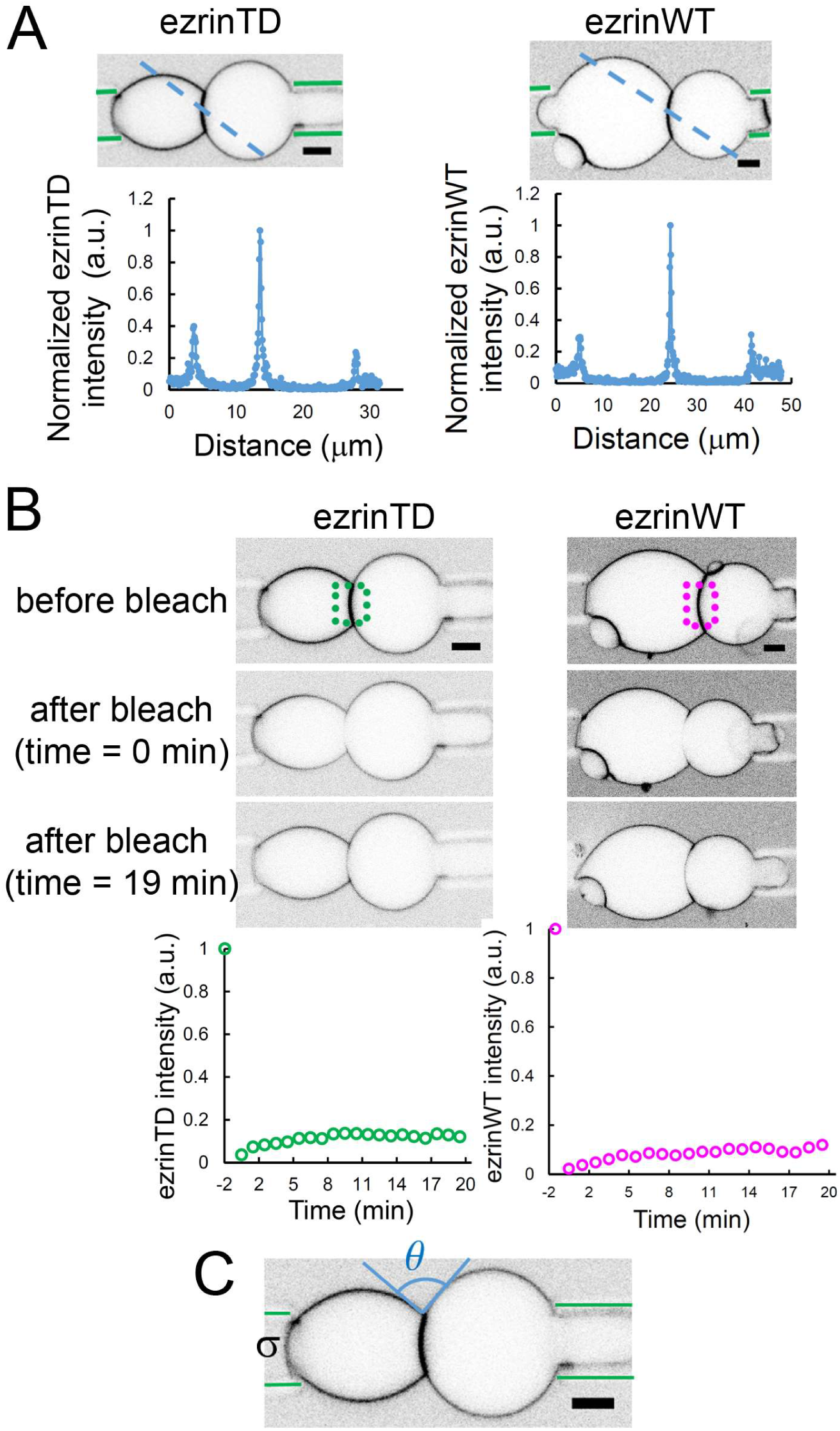
Dual micropipette experiments with GUVs tethered by ezrinTD or ezrinWT, related to Figure 1. (A) (Top) Representative confocal images of GUVs tethered by ezrinTD or ezrinWT and aspirated by micropipettes (indicated by green lines). (Bottom) Normalized ezrin fluorescence intensities along the blue dashed lines in the corresponding confocal images. Inverted grayscale images were shown. (B) (Top) Confocal images of GUVs before and after bleaching of ezrinTD or ezrin WT at the contact zone as indicated by the dotted box. Inverted grayscale images were shown. (Bottom) Fluorescence recovery after photobleaching (FRAP) experiment on ezrin-tethered membrane zones. Before photobleaching, ezrin surface fractions were 25.2% and 5.3% for ezrinTD and ezrinWT, respectively. (C) Schematic for measuring tethering strength. Scale bar, 5 μm.

**Figure S4.**
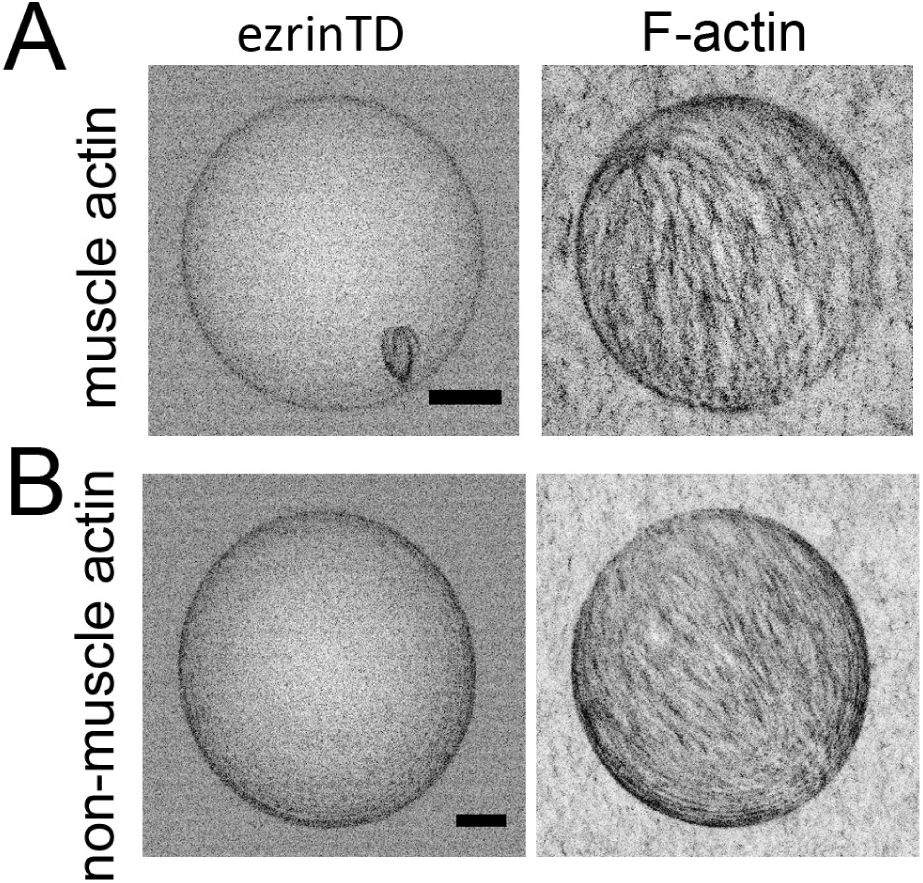
Network-like F-actin organization on ezrinTD-decorated GUV membranes, related to Figure 2. (A) and (B) Maximum intensity projects of ezrinTD decorated GUVs in the presence of F-actin in Figure 2 (A) and Figure 2 (D) were shown, respectively. Scale bars, 5 μm. Inverted grayscale images were shown.

**Figure S5.**
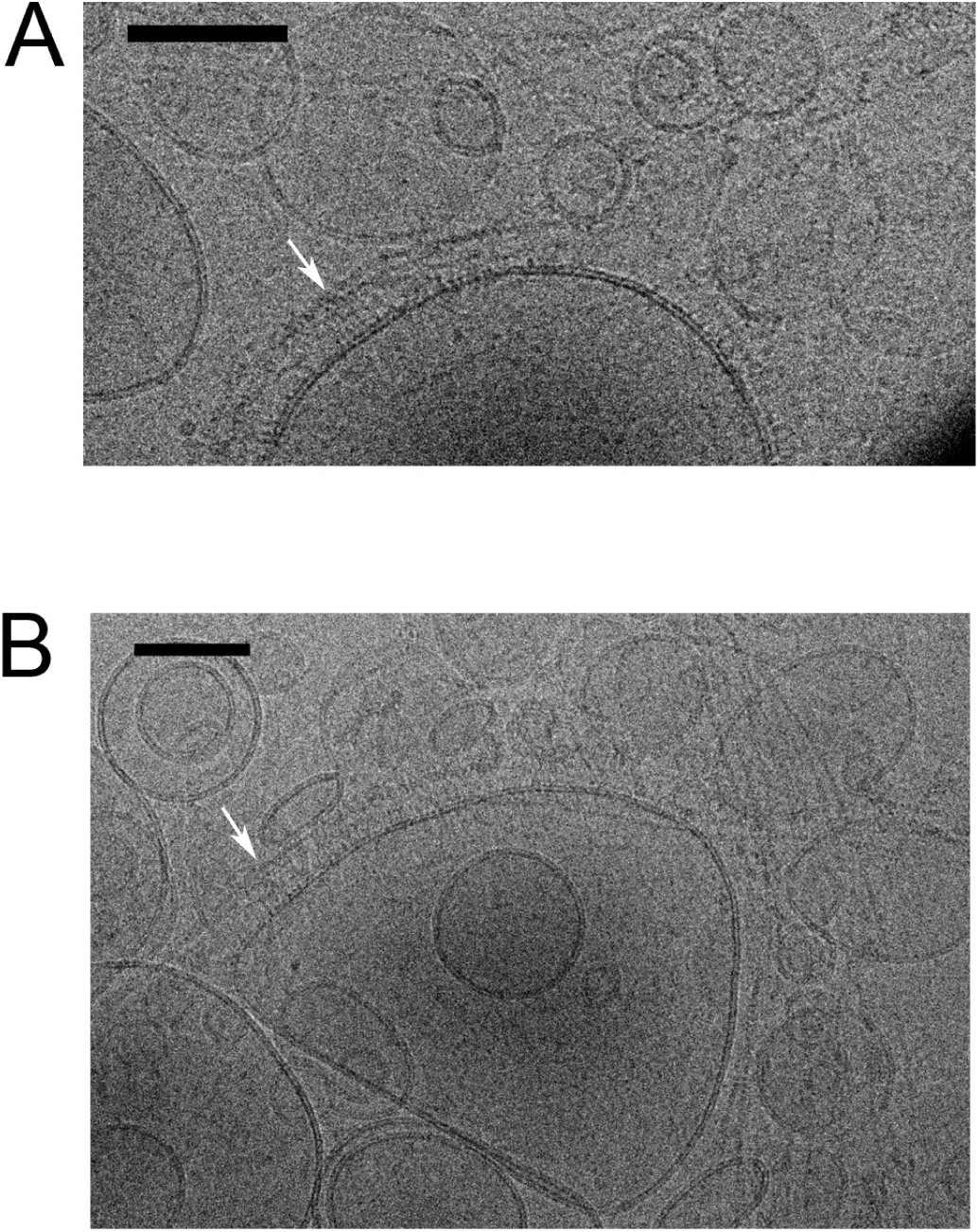
F-actin on vesicle surfaces decorated with ezrinTD, related to Figure 2. (A and B) Cryo-EM images of PIP_2_-LUVs decorated with ezrinTD and F-actin. F-actin was found to be recruited to the surface of the LUVs and followed the contour of the LUVs. Arrows indicate some F-actin along the contour of LUVs. Scale bars, 100 nm.

**Figure S6.**
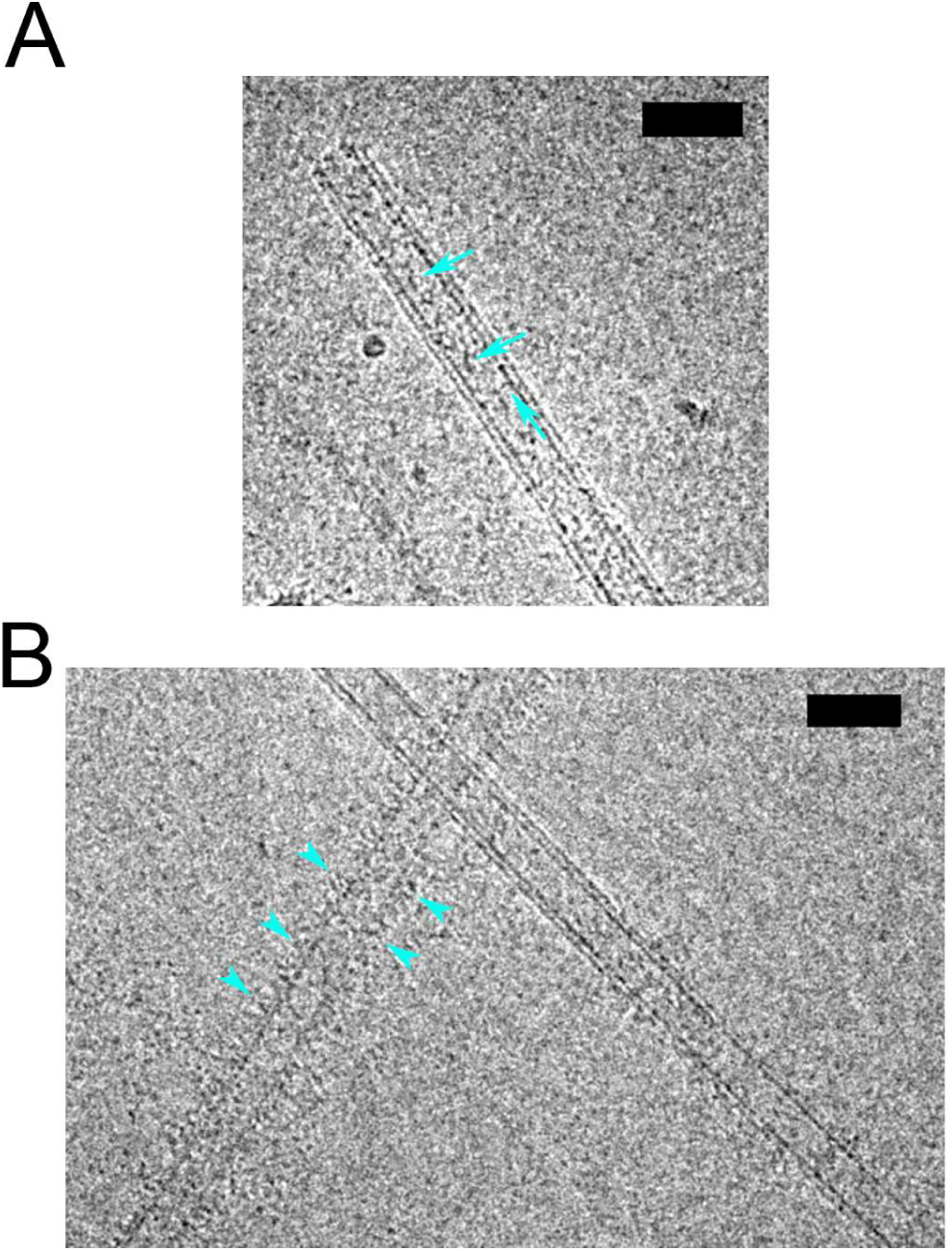
EzrinTD on isolated nanotubes, related to Figure 3. (A and B) Cryo-EM images of ezrinTD on isolated nanotubes where ezrinTD was found to either lie along the nanotubes (A), or attach perpendicularly to the nanotubes (B), assembling into brush-like organization. Arrows and arrowheads indicate some ezrinTD. Scale bars, 50 nm.

**Figure S7.**
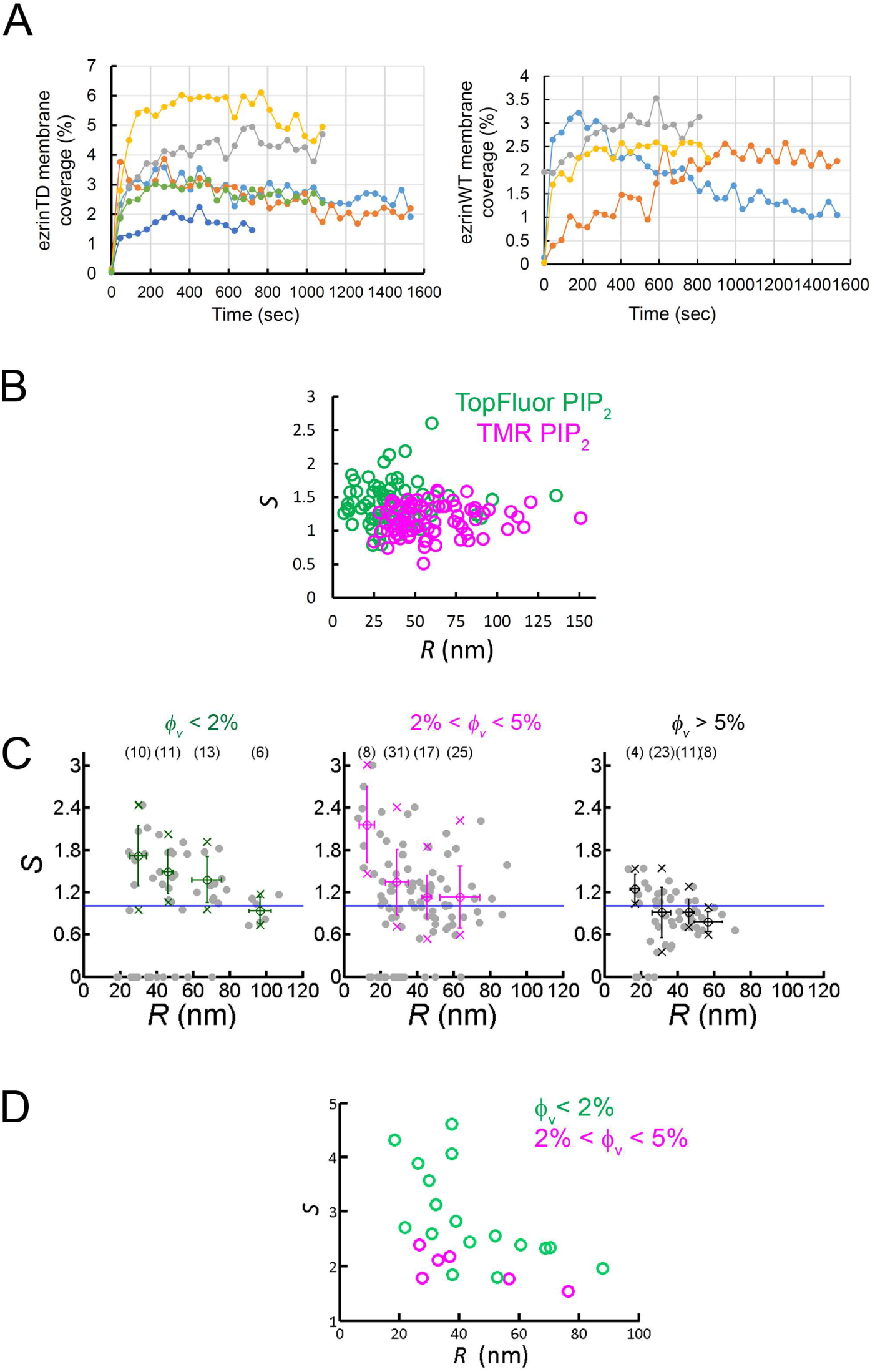
Binding of ezrin on GUVs, positive curvature-induced sorting of PIP_2_, ezrinTD and the FERM domain of ezrin on membrane nanotubes, related to Figure 3. (A) Time courses of ezrinTD and ezrinWT binding on PIP_2_-containing GUVs, showing that the steady state for protein coverage is reached in less than 200 s after protein injection. One curve represents an individual GUV measurement. (B) Absence of curvature-induced sorting of PIP_2_ measured using two types of fluorescently-labeled PIP_2_, total n =3 experiments (TopFluor PIP_2_ (N = 14 GUVs, n = 1 experiment) and TMR PIP_2_ (N = 23 GUVs, n = 2 experiments)). (C) Sorting ratio, *S*, as a function of tube radius, *R*, for ezrinTD. The same figure as in Figure 3E with circles showing individual data points. ϕ_v_: ezrin membrane surface fraction. Measurements were collected from N = 63 GUVs, sample numbers were indicated in the brackets, n > 3 independent experiments. Error bars indicate standard deviations, circles are mean and X symbols are maximum and minimum for each condition, excluding *S* = 0 data. (D) Positive curvature-induced sorting of the FERM domain of ezrin. ϕ_v_: membrane surface fraction of the FERM domain on the GUVs. *S*: sorting ratio. *R:*: membrane tube radius. Measurements were collected from N = 5 GUVs, n = 2 experiments.

**Figure S8.**
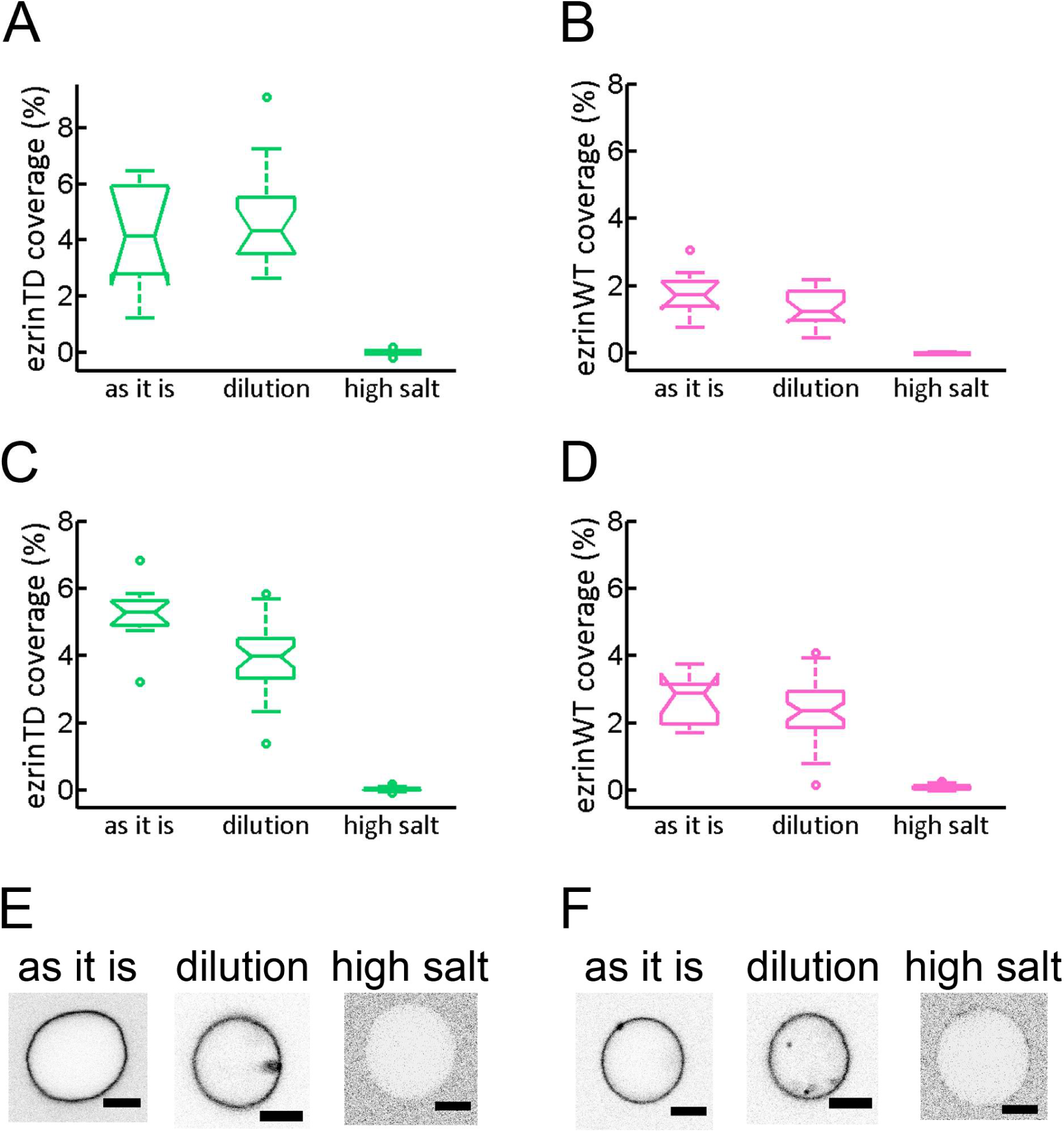
Salt-induced desorption of ezrinTD and ezrinWT bound to the outer leaflet of GUVs, related to Figure 4. (A-D) Two independent experiments of the membrane coverage of ezrinTD (A and C) and ezrinWT (B and D) at different salt conditions. GUVs were first incubated with ezrin at a concentration of 0.5 μM in a sucrose buffer (60mM NaCl, named *as it is*), followed by dilution either in a glucose buffer (60 mM NaCl, named *dilution*) or a high salt buffer 300 mM NaCl, named *high salt*) corresponding to the encapsulation experiments. Statistics: (A) N = 8, 16, 16, (B) N = 8, 18, 15, (C) N = 11, 30, 45 and (D) N = 10, 40, 40 for conditions “as it is”, “dilution” and “high salt”, respectively. Measurements were taken from distinct samples. (E and F) Representative confocal images of fluorescence signals of ezrinTD (E) and ezrinWT (F) on GUVs in the corresponding experimental conditions. Inverted grayscale images were shown. Scale bars, 5 μm.

**Figure S9.**
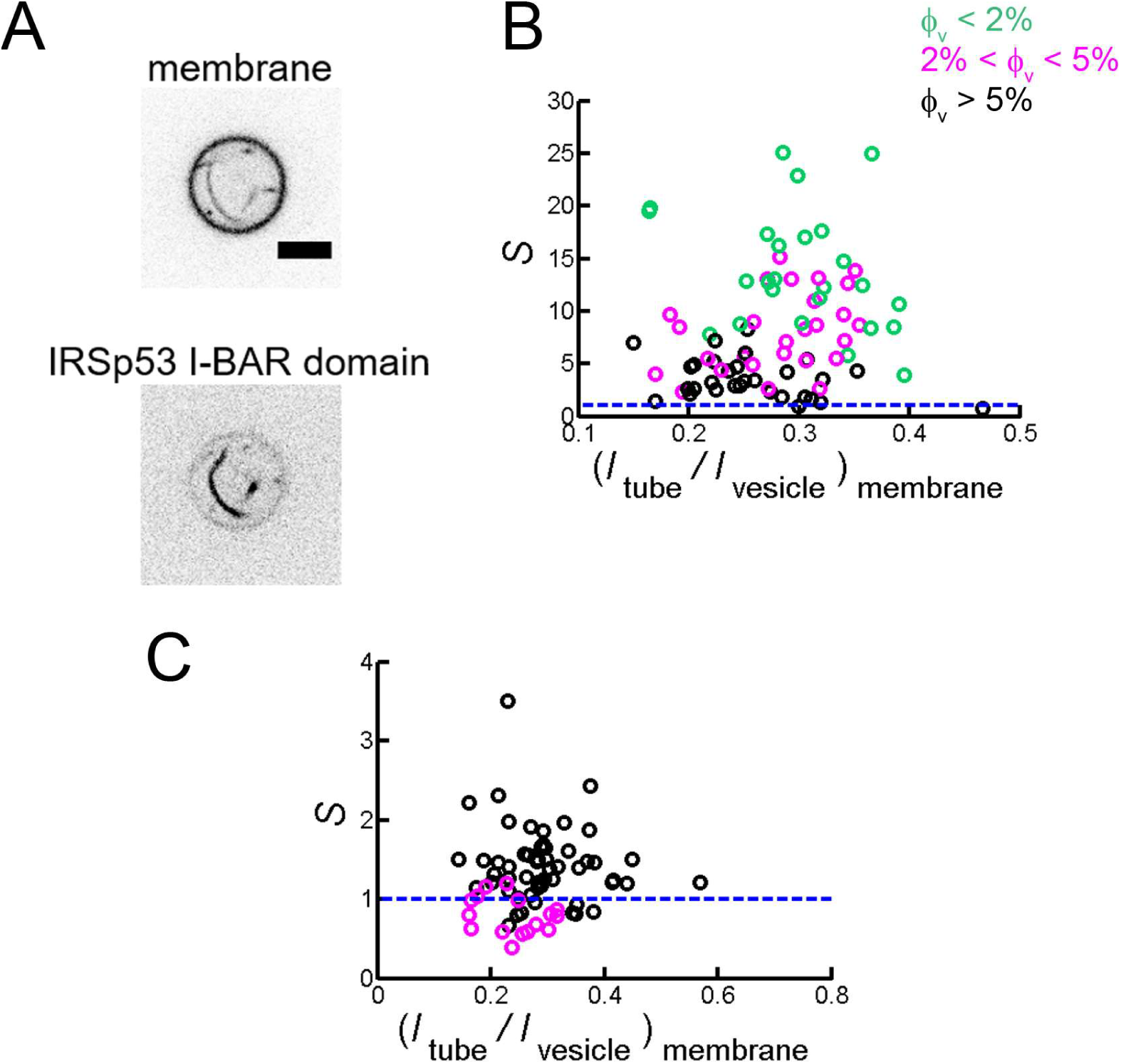
Sorting of IRSp531-BAR domain on self-induced membrane tubules and PIP_2_ enrichment in these tubules, related to Figure 6. (A) Representative confocal images of IRSp53 I-BAR domain-induced tubules. Inverted grayscale images were shown. (B) Sorting of IRSp53 I-BAR domain on self-induced membrane tubules. Maximum sorting at (*I_tube_/I_Vesicle_*)*_membrane_* ≅ 0.3 corresponds to tube radius of about 20 nm. ϕ_v_: surface fraction of the I-BAR domain on the GUVs. Measurements were collected from N = 69 GUVs, n = 2 experiments. Blue line indicates *S* = 1. (C) Sorting of PIP_2_ (black circles) and GM1* (magenta circles) in IRSp53 I-BAR domain induced tubules. I-BAR domain concentration: 0.025 μM and 0.05 μM for PIP_2_ sorting and 0.025 μM and 0.1 μM for GM1* sorting. Measurements were collected from N = 55 GUVs, n = 2 experiments for PIP_2_ sorting and N = 16 GUVs, n = 2 experiments for GM1* sorting (green circle data). Blue line indicates *S* = 1. Statistic test was performed by using *t*-test (two-sample assuming unequal variances) to compare PIP_2_ sorting and GM1* sorting, *p* = 1.2 × 10^−9^.

**Figure S10.**
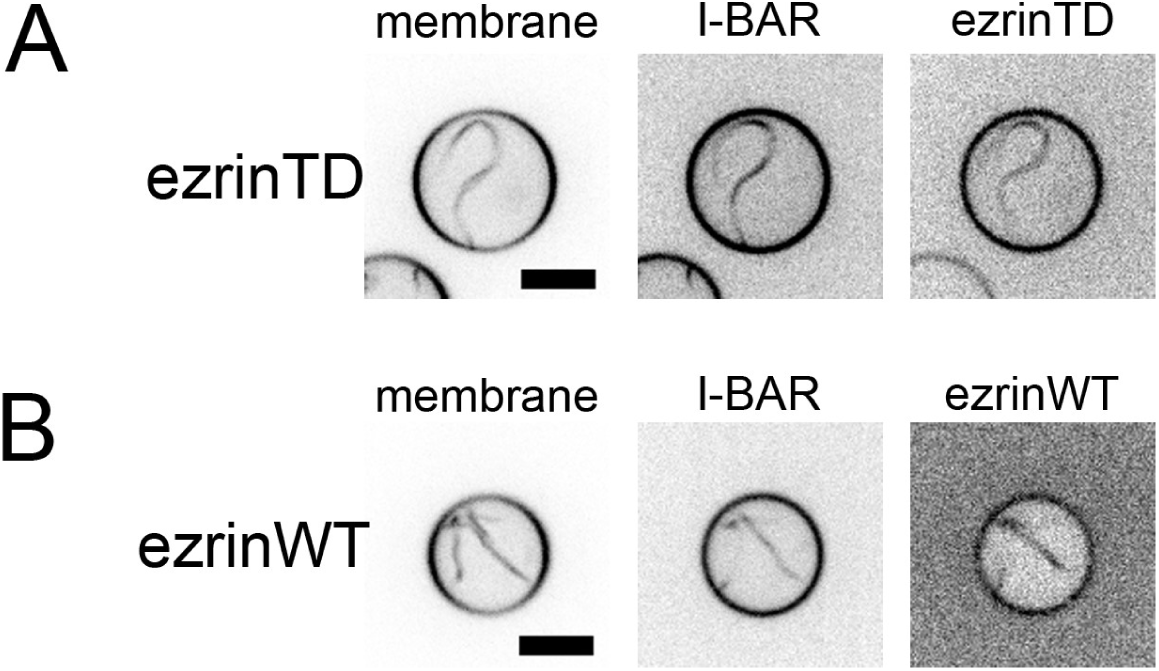
Colocalization of ezrinTD or ezrinWT with the I-BAR domain in tubules, related to Figure 6. The presence of ezrinTD (A) or ezrinWT (B) on membrane tubules induced by IRSp53 I-BAR domain. Scale bars, 5 μm. Inverted grayscale images were shown.

**Figure S11.**
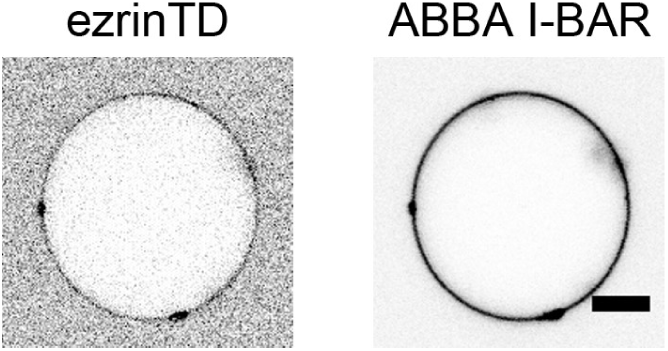
Direct interaction of ezrinTD with the I-BAR domain of ABBA, related to Figure 6. Representative confocal images of ezrinTD binding to Ni-GUVs in the absence of PIP_2_ but in the presence of ABBA I-BAR domain. Scale bars, 5 μm. Inverted grayscale images were shown.

**Video S1. Cryo-tomography of ezrinTD tethered membrane stacks, related to Figure 1.** Green colored structures represent tethered membrane stacks and magenta colored structures represent vesicular structures. Scale bars: 200 nm.

**Video S2. Cryo-tomography of ezrinTD tethered membrane stacks, related to Figure 1.** Membranes are highlighted in yellow. Scale bar: 840 nm.

